# A Hidden Binding Pocket in the β- ketoacyl-ACP Synthase FabB

**DOI:** 10.64898/2026.02.26.708327

**Authors:** Ziran Jiang, Anika Friedman, Annette Thompson, Samuel J. Andrzejewski, Kathryn Mains, Banumathi Sankaran, Michael D. Burkart, Michael R. Shirts, Jerome M. Fox

## Abstract

Assembly-line enzymes carry out multi-step synthesis of diverse metabolites by using a handful of catalytic motifs in which minor structural differences control substrate specificity and reaction order. Here we examine differences in substrate binding to FabB and FabF, the two β-ketoacyl-ACP synthases (KSs) responsible for fatty acid elongation in *Escherichia coli*, by exploring a peculiar mutational effect. In FabB, a blocking mutation in the acyl binding pocket yields a shifted, but broad product profile, while in FabF, the same mutation disrupts the binding of acyl chains longer than eight carbons (C8). X-ray crystal structures of the FabB mutant provide an explanation: a second, previously unobserved binding pocket allows medium-to-long acyl chains (≥ C8) to bind with an alternate conformation. Molecular simulations suggest that this pocket is more stable in FabB than in FabF, where mutations reduce the catalytic competency of longer chains instead of shifting them to the alternate pocket. Our findings indicate that homologous KSs differ not only in their primary binding sites but also in the availability of alternative binding modes that can buffer against mutational effects and enable functional diversification.

---

Biomolecular plasticity—the ability of proteins to adopt multiple conformations or interaction states, depending on their binding partners—enables extraordinary efficiency and adaptability in biological systems^[1,2]^. Fatty acid synthases (FASs) and polyketide synthases (PKSs) are illustrative. These assembly-line enzymes, which share a common evolutionary origin^[3]^, are found as either large multifunctional proteins (type I)^[4]^ or discrete monofunctional subunits (type II)^[5]^; both architectures build metabolites by using acyl carrier proteins (ACPs) to shuttle intermediates (“cargo”) between active sites^[6,7]^. Biomolecular plasticity allows a single ACP to bind many enzymes, and cargo-dependent differences in these interactions constrain reaction order^[8,9]^. Previous biochemical studies have resolved key details of enzyme-ACP interactions (e.g., binding surfaces, acyl binding pockets, and conformational constraints)^[8–17]^, but differences in the substrate specificities of structurally similar enzymes remain poorly understood—a major limitation for tuning the product profiles of these systems^[18,19]^.

The FAS of *Escherichia coli* (*E. coli*) is a good model system for studying enzyme-ACP interactions^[20–22]^. It initiates fatty acid synthesis with two steps: (i) FabD, an acyltransferase, transfers a malonyl group from malonyl-CoA to the phosphopantetheine (Ppant) arm of holo-ACP, and (ii) FabH, an initiating ketosynthase (KS), condenses the resulting malonyl-ACP with acetyl-CoA to form β-ketobutyryl-ACP. Subsequent rounds of enzymatic reduction (FabG and FabI), dehydration (FabZ and FabA), and elongation (FabB and FabF) yield longer acyl-ACPs (Figs. 1a and S1).

**Figure 1.**
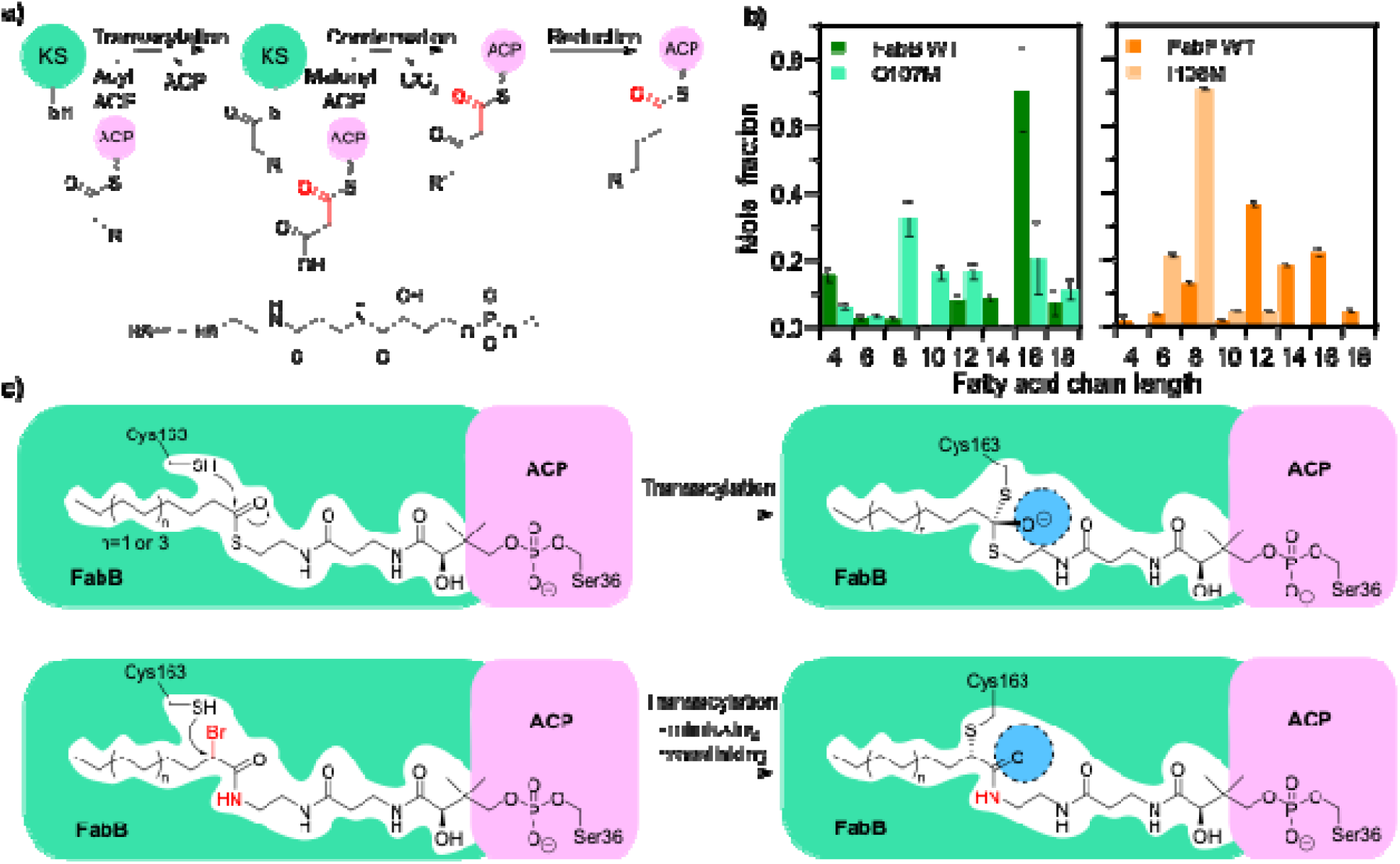
Elongating ketoacyl synthases (KSs) from *E. coli*. **a)** FabF and FabB catalyze carbon–carbon bond formation via a ping-pong mechanism: (i) an acyl-ACP transfers its acyl chain from its phosphopantetheine (Ppant) arm (below) to the catalytic cysteine of the KS to produce an acyl-enzyme, and (ii) a malonyl-ACP undergoes Clasen-like condensation with the acyl-enzyme to generate β-ketoacyl-ACP. **b)** Product profiles of an *E. coli* FAS reconstituted in vitro with either FabB or FabF as the only elongating KS (see Fig. S1 for details). Data represent the mean and standard error for n ≥ 3 technical replicates as described in the source study^[25]^. **c)** Diagrams comparing natural substrates and substrate-mimicking crosslinkers in the active sites of wild-type FabB or FabBG107M.

This study focuses on FabB and FabF, the elongating KSs at the core of fatty acid synthesis. Their substrate specificities control the pools of acyl-ACPs available for downstream enzymes and, in doing so, can shape the product profiles of oleochemical-producing pathways^[23]^. In prior work, we studied these enzymes by using the *E. coli* FAS supplemented with TesA’, a cytosolic variant of an acyl-ACP thioesterase from *E. coli*^*[23,24]*^. Within this FAS, single amino acid substitutions to the gating loop and/or acyl binding pocket of FabF and FabB shifted product profiles toward short- and medium-chain fatty acids, but the constraints imposed by specific mutations differed by KS, despite their pronounced similarity^[25]^.

Here we examine the peculiar substrate specificity conferred by the G107M mutation in FabB. This mutation sits in the acyl binding pocket, where it disrupts the binding of chains longer than eight carbons (C8). In our prior work, reconstituted (*in vitro*) FASs containing FabB_G107M_ as the only elongating KS showed a prominent increase in C8 fatty acids but still generated appreciable levels of C10-C18 (Fig. 1b)^[25]^. In contrast, the analogous mutant of FabF (FabF_I108F_) produced mainly C6-C8. X-ray crystal structures of wild-type FabB and FabF do not explain this discrepancy (Fig. S2). We speculated that FabB might have an alternative binding mode more permissive to the mutation.

We set out to examine how substrates bind to FabB_G107M_ by generating crosslinked enzyme-substrate complexes. KS-ACP interactions are weak and otherwise challenging to crystallize^[26]^. Our crosslinking strategy is based on the catalytic cycle of FabB. This enzyme has two main catalytic steps (Figs. 1a and S3)^[27]^: First, it binds to an acyl-ACP, which transfers an acyl chain to Cys163 to form an acyl-enzyme intermediate. Second, it binds to malonyl-ACP, which displaces holo-ACP and undergoes Claisen-like condensation with the acyl-enzyme to generate a β-ketoacyl-ACP. We used ACPs with α-bromopantetheine amide and chloroacrylate to trap the transacylation and condensation states, respectively (Figs. 1c and S3-S5)^[27]^. For α-bromopantetheine amide, we used crosslinkers **1-6**, which mimic C2, C4, C6, C8, C12, and C16 substrates; for chloroacrylate, we used **7** and **8**, which mimic C8 and C14 substrates (Fig. S4)^[27]^. In initial crosslinking experiments, FabB_G107M_ showed a clear preference for C8 (i.e., **4** and **7**), consistent with its product profile; yields for other substrate lengths were modest (Fig. S6).

We focused X-ray crystallography on C8 and C12 substrate mimics, which we expected to have different sensitivities to the G107M mutation. We optimized crystallization conditions for crosslinkers **4** and **5** (C8 and C12 acyl chains), which afforded good crosslinking efficiencies; the resulting crystal structures had resolutions of 2.14 Å and 2.25 Å, respectively (Table S1). In our structures, FabB_G107M_ forms a homodimer crosslinked with two symmetrically positioned ACPs (Fig. 2a). The enzyme-ACP interfaces are comparable between the two complexes with subtle differences. For ACP_C8_, acidic residues Glu47, Glu48, and Asp56 on helix 1 form salt bridges with Lys124, Lys127, and Arg45 on FabB, while Gly33, Asp35, and Asp38 on loop 1 of ACP form additional interactions with Lys63 and Arg66 on FabB (Figs. 2b-d). For ACP_C12_, Met44 (backbone) and Glu47 on helix 1 form polar salt bridges with Arg124 and Lys127, respectively; and Gln14, Gly33, and Asp38 on loop 1 of ACP form hydrogen bonds (h-bonds) with Arg62, Lys63, Arg66, and Asp35 on FabB (Figs. 2e-g). These interfaces are consistent with previous structures of wild-type FabB in complex with crosslinked ACP substrates^[28]^, indicating native protein-protein interactions.

**Figure 2.**
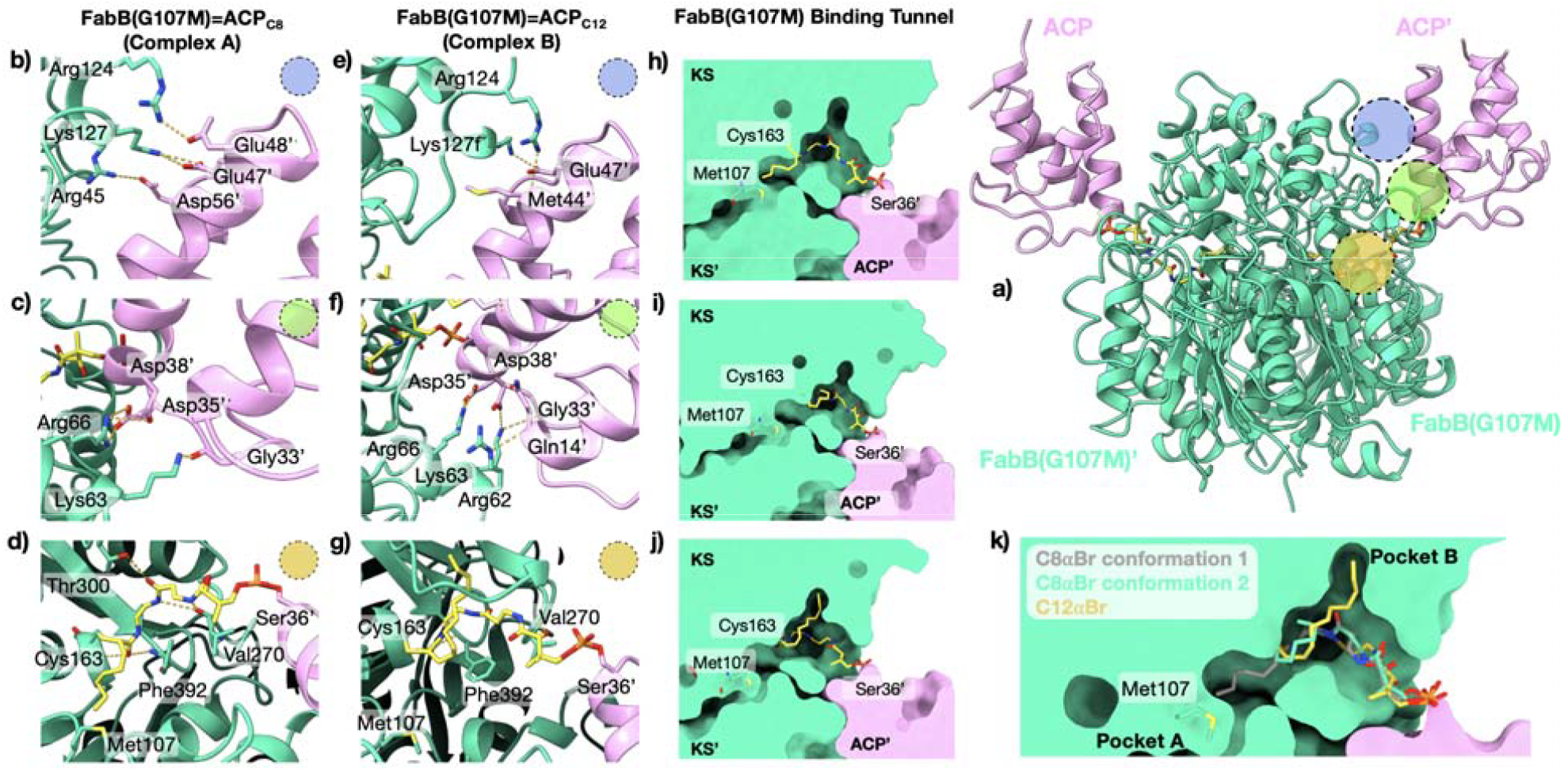
X-ray crystal structures of FabB_G107M_ show substrate-specific binding interactions. **a)** The X-ray crystal structure of FabB_G107M_=ACP_C8_ (PDB entry 9MLW) shows a homodimer of FabB with two symmetrically positioned ACPs. **b-d)** The FabB_G107M_=ACP_C8_ shows key interactions between FabB and **b)** helix 1, **c)** loop 1, or **d)** the Ppant arm of ACP. **e-g)** The FabB_G107M_=ACP_C12_ (PDB entry 9MLX), shows key interactions between FabB_G107M_ and **e)** helix 1, **f)** loop 1, or **g)** the Ppant arm of ACP. **h-j)** In the FabB_G107M_=ACP_C8_, the acyl chain **h)** extends into pocket A or **i)** folds back to the entrance of pocket B. **j)** In the FabB_G107M_=ACP_C12_ complex, the acyl chain shares the second conformation and enters pocket B. Colored circles highlight corresponding regions of the main structure. **k)** Overlay of C8 and C12 crosslinkers from FabB_G107M_=ACP_C8_ and FabB_G107M_=ACP_C12_ demonstrates the acyl substrate chain length-based conformational changes point to pockets A and B.

The two crystal structures show similar arrangements of the substrate with important differences in acyl chain positioning. For both chain lengths, covalent bonds between (i) the pantetheine arm and Ser36 (ACP) and (ii) the crosslinker α-carbon and Cys163 have partial density, and the amide carbonyl sits in the oxyanion hole formed by the backbone amides of Cys163 and Phe392. In FabB_G107M_=ACP_C8_, the acyl chain adopts two conformations (Figs 2h-i): In the first, the C8 carbon sits near Met107 in the truncated tunnel, hereafter referred to as “pocket A”. In the second, the acyl chain bends away from this tunnel. The polder map (σ=2.5) has density for both conformations (Fig. S7). For FabB_G107M_=ACP_C12_, the acyl chain adopts only the second conformation, where the C6-C12 carbons define a novel pocket formed by His298-Thr302 and Gly305-Val307 on FabB, hereafter referred to as “pocket B” (Fig. 2j). The polder map shows no density in pocket A (Fig. S7). Pocket B may explain the surprising activity of FabB_G107M_ on acyl chains of eight carbons or longer.

We examined the influence of G107M on access to pockets A and B using molecular dynamics (MD) simulations combined with a recently developed enhanced sampling method, termed multiple topology replica exchange of expanded ensembles (MT-REXEE)^[29]^. MT-REXEE allows acyl chains to grow and shrink during the simulation, in this case in two-carbon increments (C2 to C4, C4 to C6, etc., from C2-C16 acyl chains). This feature accelerates access to alternative binding modes and enables estimates of relative binding free energies between alternate pockets as a function of chain length. We examined both non-covalent enzyme-ACP complexes and covalent acyl-enzyme intermediates for FabB_WT_, FabB_G107M_, FabF_WT_, and FabF_I108M_. For each state and each enzyme, we ran three replicate simulations—two with the acyl chain initiated in pocket A and one with it initiated in pocket B—and tracked catalytically competent complex conformations, here defined as the distance between Cys163 and the catalytic carbonyl less than 4 Å.

Our simulations showed three pockets that allow the acyl chain to adopt a catalytically competent conformation: A and B, as defined above, and C, where the acyl chain is parallel to the Ppant arm (Fig. 3a). Figures 3b-3e show the percent of each total enzyme-ACP simulation in which the acyl chain samples pockets A-C (defined by RMSD to a reference structure; see Fig S8). In both FabB_WT_ and FabF_WT_, chains show a clear preference for pocket A (40-70% of each trajectory, depending on chain length). Given the observation that the FabB_G107M_ variant maintains activity on substrates longer than 8 carbons, but the analogous FabF_I108F_ does not (Fig. 1b), we hypothesized that pocket B might be accessible in FabB but not in FabF^[30]^. Indeed, sampling of pocket B is 3-to 10-fold higher in FabB than in FabF, indicating that it is more accessible (Fig S9). The G107M mutation in FabB shifts the binding preference of chains longer than eight carbons from A to B, whereas the I108M mutation in FabF causes these chains to spend a larger fraction of the trajectory in a non-catalytically competent conformation (i.e., the acyl chain oriented away from catalytic residues). Simulations of the covalent acyl-enzyme intermediate show similar trends (Fig. S10). The MT-REXEE analyses provide strong support that pocket B is not an artifact of X-ray crystallography or covalent crosslinking, but rather an alternative, functionally relevant pocket in FabB.

**Figure 3.**
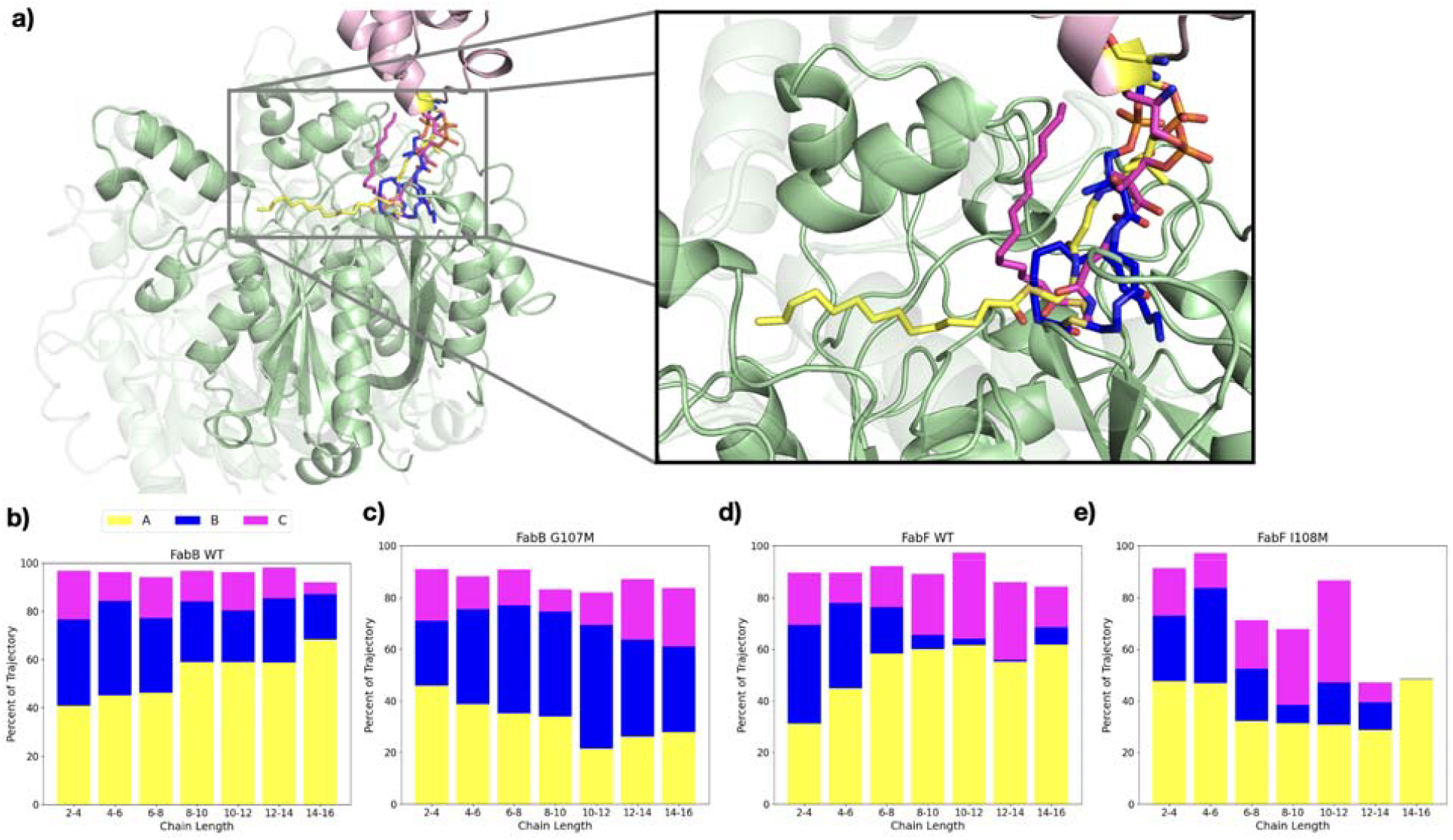
Molecular dynamics (MD) simulations of elongating KSs. **a)** Centroid structures from MD simulations of FabB bound to ACP_C12_ show interactions with pockets A (yellow), B (blue), and C (pink). **b-e)** These plots show the percent of the MT-REXEE trajectory in which the acyl chains sample each pocket with a catalytically competent conformation (a distance between C163 and the catalytic carbonyl less than 4 Å). The full color bar represents the mean percentage of all trajectories for which the complex was in a catalytically competent state, and the color divisions denote which pocket the acyl chain occupied in that catalytically competent state. **b-c)** For FabB, the G107M mutation reduces sampling of pocket A and favors pocket B. **d-e)** For FabF, I108M does not cause a prominent increase in sampling of pocket B but instead destabilizes the enzyme-substrate complex, as indicated by a significant decrease in the overall catalytic competency of the pocket. Data represent the mean of n = 3 simulations for each chain length with error described in Fig. S9.

MT-REXEE also enables estimates of the energetic penalties that mutations impose on binding. In simulations of FabB-ACP complexes, G107M imposes a penalty of 2-3 kcal/mol on the extension of C6 to C8, C8 to C10, and C10 to C12 within pocket A (Fig. S11); I108M imposes the same penalty for FabF. In the covalent intermediate, both mutants yield a smaller extension penalty of 1 kcal/mol for the C6 to C8 transformation in pocket A, which may reflect the enhanced flexibility of surrounding loops in the absence of ACP (Fig. S12). Penalties for extensions immediately beyond C8 (i.e., C8 to C14), in turn, were negligible or very small. When we carried out a similar set of analyses for pocket B, we observed no significant free energy differences between mutant and wildtype for extensions from C6 to C10 (Figs. S11-S12). For C10 to C12, the mutant was destabilizing in the FabF-ACP complex and modestly stabilizing for FaB-ACP, but it destabilized this extension only very slightly for both covalent intermediates—overall, a modest effect. Altogether, our results suggest that G107M increases sampling of pocket B primarily by destabilizing binding to pocket A.

The enhanced accessibility of pocket B in FabB is surprising, given the structural similarity of FabF. We used our simulations to explore the molecular basis of this difference. Across simulations of WT and mutants, interactions between FabB or FabF and the ACP interface do not show major differences, regardless of the occupied pocket (A or B; Fig. S15); however, the interaction networks that stabilize the Ppant linker when the acyl chain occupies pocket B are distinct. In FabB, Met206 and Lys310 form h-bonds with the Ppant linker, but in FabF, the analogous residues Ala204 and Ala312 do not (Fig. S16, S17A-B). Val272 and Ala273 also form weak transient interactions with the Ppant linker when the acyl chain occupies pocket B of FabB, while in FabF, the corresponding residues Thr270 and Ser271 form h-bonds that may limit linker flexibility (Fig. S16, S17B-C). Ultimately, our simulations suggest that differences in the accessibility of pocket B between the two KSs result from differences in minor stabilizing effects, rather than major structural obstructions.

Structural overlays of FabB and FabF bound to various substrates and substrate analogues provide context for the role of pocket B in the catalytic cycle of KSs (Fig. S18a-b). Previous structures of FabF_WT_=ACP_C8_ and FabF_WT_=ACP_C16_ (PDB entries 6OKG and 7L4L) suggest that acyl chain loading is regulated by a gating mechanism in which ACP binding shifts Phe400 from closed to open, yielding access to the active site (Figs. S18b and S18f). Although no crystal structures of FabB include this conformation, our MD simulations show that Phe392 transitions between “gate-open” and “gate-closed conformations (Fig. S13). A combination of these previous studies suggests the following catalytic cycle (Fig. S19): (i) FabB and FabF begin in “gate-closed” conformations (Fig. S19a). (ii) ACP binding induces a “gate-open” conformation in which conserved histidine residues stabilize the thioester carbonyl and enable exchange of the acyl chain between pockets A, B, and C (Fig. 15b). (iii) Closure of the Phe gate blocks access to pocket C and restores an oxyanion hole that stabilizes the tetrahedral intermediate required for acyl chain transfer. (iv) When transfer is complete, holo-ACP unbinds from the enzyme, leaving a stable acyl-enzyme intermediate (the starting point for subsequent condensation with malonyl-ACP).

The proposed catalytic steps suggest distinct roles for each pocket in FabB. Pocket C is a transient binding site that enables acyl chain delivery to the active site. Pocket A is the primary binding tunnel during acyl chain elongation. And pocket B may help tune substrate specificity. Our prior work on FabB suggests a mechanism of negative cooperativity that is absent in FabF ^[31]^; It is evident in the crystal structure of FabB_WT_= ACP_C16_ (Fig. 4). In the FabB homodimer, binding of a long acyl chain (C14-C16) to pocket A of one monomer causes a gate defined by Glu200-Gln113’—hereafter, the E-Q gate—to open; in response, the symmetric gate on the other subunit (Glu200’-Gln113) closes and inhibits the binding of chains longer than twelve carbons to pocket A’. By providing an alternative binding mode, pocket B may alleviate—though not completely eliminate—this autoinhibition. This theory of substrate specificity is consistent with the product profiles measured in our *in vitro* assays, which suggest that FabB_WT_ is less active on C16 than FabF_WT_ yet more accommodating of an obstruction in pocket A (Fig. 1b).

**Figure 4.**
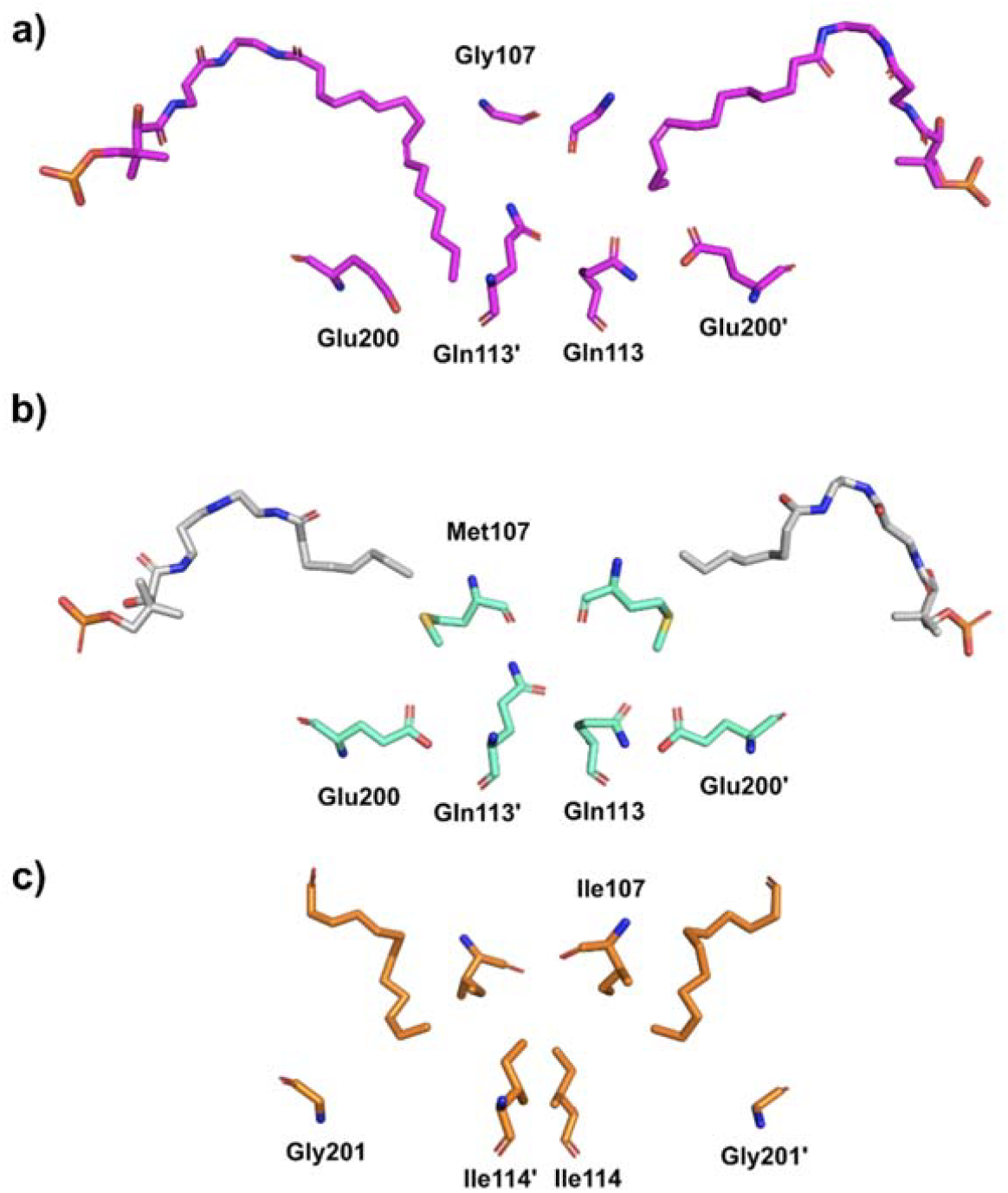
Negative cooperativity within the enzyme homodimer governed by a back gate in FabB (E-Q gate). a) In FabB_wt_=ACP_C16_ (PDB entry 72Z9), acyl chain binding to the left pocket causes Glu200 to shift out, while Glu200 and Gln113’ to adopt an open conformation that pushes Gln 113 and Glu200’ into a closed conformation, hindering substrate binding to the right pocket. Note: The last three carbons of the acyl tail in the right pocket are not resolvable in this structure. b) In FabB_G107M_=ACP_C8_ (PDB entry 9MLW), Gly107Met prevents both acyl chains from reaching the bottom of the acyl binding pocket, and the E-Q gate is not activated (i.e., Glu200 and Glu 200’ are closed). c) A crystal structure of FabF covalently linked with dodecanoic acid (PDB entry 2GFY) shows a broad acyl binding pocket with no E-Q gate.

Assembly-line enzymes use seemingly endless variations on a small set of catalytic motifs to carry out multi-step syntheses. Our analysis suggests that features as simple as an acyl binding pocket can obscure functionally relevant structural differences that govern substrate routing. We find that homologous KSs diverge not simply in substrate preference, where differences are minor for the WT enzymes, but in the availability of alternative, energetically accessible binding modes that buffer against mutational perturbations. Follow-up analyses of FabB binding to cis-dec-3-enoyl-ACP, on which FabB is markedly more active than FabF, could clarify whether the newly discovered pocket contributes to unsaturated fatty acid biosynthesis. Beyond KSs, our findings illustrate how hidden conformational landscapes can drive functional diversification in enzyme families central to metabolism.

## SUPPORTING INFORMATION

The authors have cited additional references within the Supporting Information.

## ACKNOWLEDGEMENTS

This work was supported by the DOE Office of Science, Bio- logical and Environmental Research Program (A.J.F., A.T., S.J.A., and J.M.F., DE-SC0023142), the United States Army Research Office (K.M., award W911NF-18-1-0159), the Howard Hughes Medical Institute, and the National Institutes of Health, the National Institute of General Medical Sciences (B.S., ALS-ENABLE grant P30 GM124169; Z.J. and M.D.B., R01 GM095970). The Advanced Light Source is a Department of Energy Office of Science User Facility under contract no. DE-AC02-05CH11231.

## FIGURES

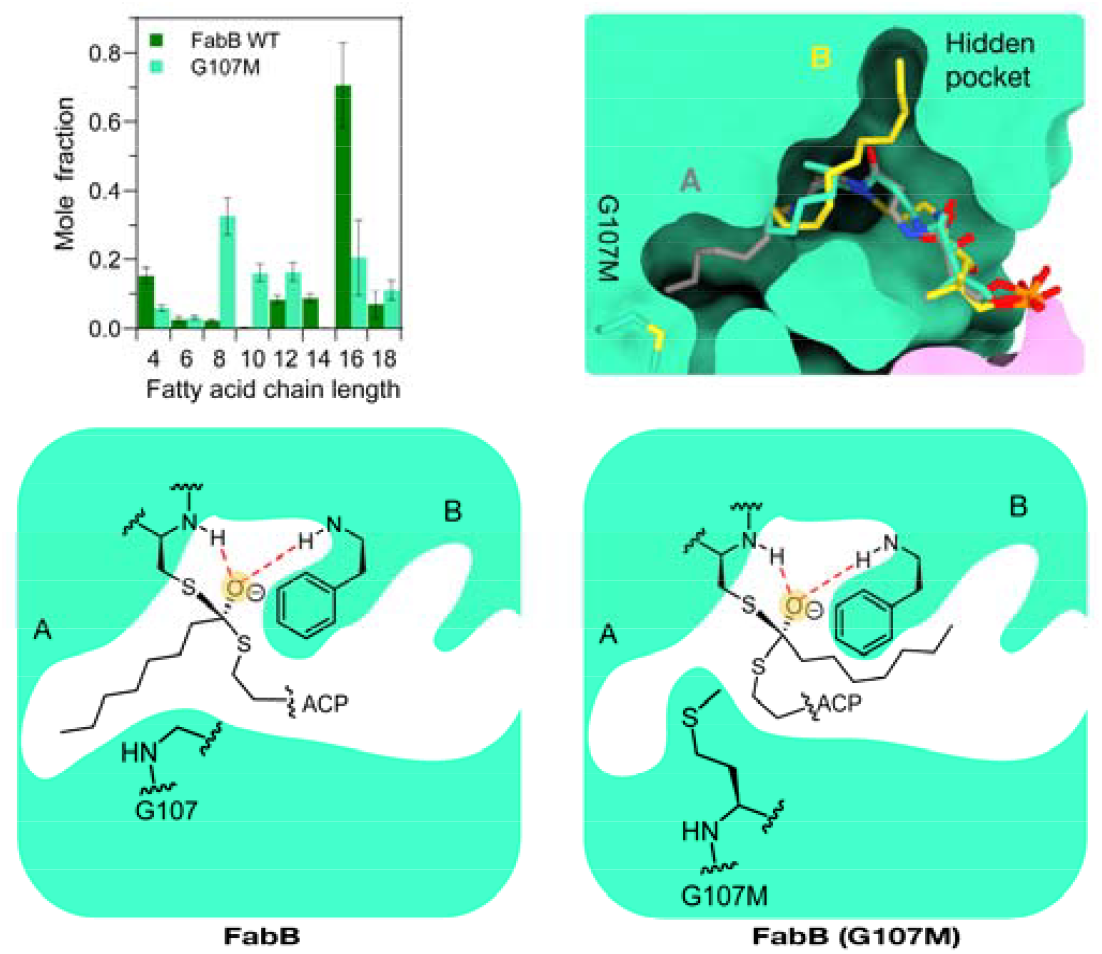

**Table of Contents**. Homologous KSs diverge not only by substrate preference, but also in the availability of alternative, energetically accessible binding modes. In this study, we examine FabB, a β-ketoacyl-ACP synthase from *Escherichia coli*. Its unexpected tolerance of the G107M mutation in (A) the canonical binding pocket led us to uncover (B) a secondary pocket that buffers against mutational perturbations.

