## Supporting Information for "A Hidden Binding Pocket in the β- ketoacyl-ACP Synthase FabB"

### SI METHODS

**General experimental methods, materials, and reagents.** We used NEB® Stable Competent E. coli and BL21(DE3) E. coli cells (New England Biolabs) for cloning and protein expression, respectively. We obtained isopropyl β- d-1-thiogalactopyranoside (IPTG) from Thermo Fisher Scientific and arabinose from Acros Organics. antibiotics from Thermo Fisher Scientific (carbenicillin and kanamycin sulfate) and Millipore Sigma (chloramphenicol); and media components from Thermo Fisher Scientific (tryptone, yeast extract, and S.O.C media). For buffer preparation, Tris base and HCl, DTT, and DMSO were purchased from Fisher Scientific. $MgCl_2$ was obtained from BDH Chemicals, and NaCl was obtained from RPI Research Products. For protein purification, we purchased all chromatography columns, including Hiload Superdex 200 pg (16 × 600), HiLoad Superdex 75 pg (16 × 600) size exclusion columns, and an FPLC (ÄKTA Pure) from Cytiva. We synthesized site-specific crosslinkers using the previously developed protocol with purity >98% by NMR. We prepared fresh stock solutions of crosslinker in DMSO for each experiment. For electrophoresis gel preparation and evaluation, we used SDS (Sigma Aldrich), 40% acrylamide/bis-Acrylamide 29:1 (Fisher Scientific), TEMED (Sigma Aldrich), and APS (Fisher Scientific). For SDS-PAGE gel staining and imaging, we used Coomassie Brilliant Blue staining and Perfection V19 scanner (Epson). For X-ray crystallography, we purchased crystallization reagents from Hampton Research and cryoprotectants from MiTeGen.

**Plasmid construction.** Table S2 describes the details and sources of all plasmids used in this study.

**Expression and purification of *E. coli* FAS proteins, including FabF mutants and FabB mutants.** We overexpressed all FAS enzymes in *E. coli* as described previously[1,2]. In brief, we transformed chemically competent BL21(DE3) cells with individual plasmids, recovered the cells in S.O.C. (Super Optimal broth with Catabolite repression) media (2% tryptone, 0.5% yeast extract, 10 mM NaCl, 2.5 mM KCl, 10 mM $MgCl_2$, 10 mM $MgSO_4$, and 20 mM glucose) for an hour, plated them on Luria–Bertani (LB) agar plates supplemented with antibiotic (10 g/L tryptone, 5 g/L yeast extract, 10 g/L sodium chloride; 50 µg/mL carbenicillin or kanamycin), and grew the cells at 37ºC overnight. Picking individual colonies, we inoculated 20 mL cultures of LB media containing antibiotic (50 µg/mL carbenicillin or kanamycin) and incubated them at 37 ºC and 225 rpm until the cultures became cloudy (~5 hours or overnight). We diluted the 20-mL cloudy cultures into 1 L of rich induction media supplemented with antibiotics (20 g tryptone, 10 g yeast extract, 5 g sodium chloride, 0.4% glucose, 72 mL of 5X M9 solution; 50 µg/mL carbenicillin or kanamycin) and grew the 1-L cultures at 37 ºC and 225 rpm until $OD_{600}$ reached 0.4-0.8. At this OD, which is indicative of mid-exponential growth, we added 0.5 mM IPTG and transferred the cultures to 22 ºC and 225 rpm for another 14-18 hours. We harvested cells by centrifuging them at 5,000 rpm for 15 minutes, and we froze cell pellets at -80 ºC for future use. We purified proteins using a fast protein liquid chromatograph (FPLC, AKTA Pure). We lysed cell pellets by adding a standard lysis buffer: 4 mL of buffer (B-PER) with 2 mg TAME, 2 mg magnesium sulfate heptahydrate, 3.5 mg TCEP, 3.75 µl PMSF, 0.5 mg lysozyme, and 200 units DNase I (New England Biolabs) per g of cell pellet. We isolated histidine-tagged protein by flowing lysate over a nickel-affinity column (HisTrap HP column). For standard purifications, we used 50 mM Tris-HCl, pH 7.5, 0.5 mM TCEP, 300 mM NaCl, and 0-500 mM imidazole. For all proteins, we buffer-exchanged isolated protein with a spin concentrator (3-10kDa VivaSpin; Sartorius) and purified it further, where necessary, with an anion-exchange column (HiPrep Q HP 16/10 column with 50 mM Tris-HCl, pH 7.5 or 8.5, 0.5 mM TCEP, and 0-1 M NaCl) or a cation-exchange column (HiPrep SP HP 16/10 column with 50mM sodium phosphate, 0.5 mM TCEP, 20% glycerol, and 0-1 M NaCl). We used standard procedures to confirm and store purified proteins. We confirmed protein purity and molecular weight by using sodium dodecyl sulfate polyacrylamide gel electrophoresis (SDS-PAGE), carried out a final buffer exchange, and concentrated purified proteins with a spin concentrator (5 or 10 kDa VivaSpin as before). For all proteins, we measured final concentrations with a Bradford assay by using bovine serum albumin (BSA) as a standard. We flash froze purified proteins and stored them in 20% glycerol at -80 ºC. For proteins of low abundance, we used only nickel-affinity chromatography, which was typically sufficient to achieve single-band purity with SDS-PAGE (e.g., E. coli FabZ).

**Expression and purification of *E. coli* acyl carrier protein ACP.** We followed our previous protocol to express and purify ACP[3]. Briefly, we synthesized the gene of native *E. coli* ACP by Twist Bioscience with a C-terminal stop codon to circumvent the expression of the C-terminal 6xHis-tag on the vector. It was cloned into a pET-21 vector containing NdeI and XhoI restriction sites. We transformed the ACP gene into *E. coli* BL21(DE3) cells and cultured them in 8 mL of LB broth (LB broth Miller, RPI International) prepared with 25 g/L in deionized water, followed by autoclaving. After adding ampicillin (100 mg/L) and inoculation, we incubated the starter culture on a spinning wheel for 12 h at 37 ºC. We added the starter culture into LB broth (1L) with 100 mg/L of ampicillin and incubated at 37 ºC with 150 rpm shaking until the $OD_{600}$ reached between 0.6 and 0.8. We then induced the culture by adding 0.5 mM IPTG at 18 ºC. After shaking for 12 h at 18 ºC, we collected cell pellets through centrifugation at 2,000 rpm for 30 min and stored them at -20 ºC or used them directly for lysis in the next step. For lysis, we resuspended the cell pellets with cold 50 mM Tris-HCl buffer, pH 7.4, 150 mM NaCl, 10% glycerol. We then sonicated the resuspended pellets on ice for 10 min with 1 s on and 3 s off cycles. We removed the insoluble impurities by centrifugation at 4,000 rpm for 1 h. We subjected the supernatant to isopropanol titration with an equal volume of isopropanol. We removed insoluble impurities by 5,000 rpm for 1 h to obtain the ACP mixture in 50% aq. isopropanol. We directly loaded the supernatant onto a HiTrap Q HP anion exchange column and purified *via* FPLC (ÄKTA Pure) with a linear gradient of 0 to 1M NaCl in 50 mM Tris-HCl buffer pH 7.4. We evaluated eluted fractions by SDS-PAGE (12% acrylamide). We pooled and dialyzed the pure ACP fractions using 3.5K MWCO SnakeSkin dialysis tubing (Thermo Fisher Scientific) with 50 mM Tris-HCl buffer pH 7.4 and 150 mM NaCl. We spin concentrated these samples to 5 mg/mL using spin concentrator (Millipore Amicon Ultra) with 3K MWCO. We aliquoted, flash-froze, and then stored the concentrated samples at -80 ºC or used them directly.

**Expression and purification of acyl carrier protein phosphodiesterase (AcpH).** We expressed AcpH with an N-terminal 6xHis-tag in *E. coli* BL21(DE3) cells, following our previous method[3]. We prepared starter culture with the inoculation of a single colony in LB broth (5 mL) with 50 mg/L kanamycin for 12 h at 37 ºC. We then added the culture into LB broth (1L) with 50 mg/L kanamycin and allowed it to grow at 37 ºC with shaking at 150 rpm to reach an $OD_{600}$ of 0.6. We induced the culture with 1 mM IPTG at 16 ºC. We incubated the induced culture for 12 h at 16 ºC with shaking at 120 rpm. We collected the cell pellets by centrifugation for 30 min at 2,000 rpm. With cold lysis buffer 50 mM Tris-HCl, pH 7.5, 500 mM NaCl, 10% glycerol, we resuspended the cell pellet and sonicated it on ice for 10 min with 1s on and 3s off cycles. We performed sequential centrifugation at 5,000 rpm for 1 h to remove insoluble impurities, resulting in a clear supernatant. We purified the supernatant using Ni-NTA resin (Thermo Fisher Scientific) at 4 ºC in a cold room. We washed the column with 10 column volumes (CV) of the aforementioned lysis buffer, followed by another wash with lysis buffer containing 10 CV of 10 mM imidazole. We used lysis buffer containing 250 mM imidazole to elute AcpH. We dialyzed the eluted AcpH using 10K MWCO SnakeSkin dialysis tubing (Thermo Fisher Scientific) in lysis buffer for 12 h at 4 ºC. Using an Amincon Ultra spin concentrator (Millipore) with a 10 kDa MW cutoff, we concentrated the dialyzed protein to 5 mg/mL and flash-froze it in aliquots for storage at -80 ºC or used it directly.

**Expression and purification of 4'-phosphopantetheinyl transferase (Sfp).** We recombinantly expressed Sfp with an N-terminal 6xHis-tag in *E. coli* BL21(DE3), as discussed previously[3]. We grew single colonies in LB broth (5 mL) with 100 mg/L ampicillin for 12 h at 37 ºC to yield a starter culture. We inoculated LB broth (1L) with 100 mg/L ampicillin using the starter culture and incubated at 37 ºC with shaking at 150 rpm until an $OD_{600}$ of 0.7 was reached. We induced the resulting cultures with 0.5 mM IPTG at 18 ºC. We allowed the induced culture to grow at 18 ºC for 16 h, then centrifuged at 2,000 for 30 min to obtain cell pellets. We resuspended the cell pellets in cold lysis buffer (50 mM potassium phosphate, pH 7.5, 150 mM NaCl, 10% glycerol) and lysed by sonication (1 s on, 3 s off cycles for 10 min) on ice. We conducted centrifugation at 5,000 rpm for 1 h to remove insoluble impurities. We loaded the resulting supernatant onto a Ni-NTA column. We washed the column with 10 CV of lysis buffer, followed by 10 CV of lysis buffer containing 10 mM imidazole. We eluted 6xHis-tagged Sfp with lysis buffer containing 250 mM imidazole. We performed sequential dialysis via 10K MWCO SnakeSkin dialysis tubing (Thermo Fisher Scientific) in lysis buffer (1L) for 12 h at 4 ºC. Using a 10K MWCO Amicon Ultra spin concentrator (Millipore), we concentrated the dialyzed protein to 5 mg/mL and flash-froze it in aliquots for storage at -80 ºC or used it directly in the next step.

**Expression and purification of CoaA, CoaD, and CoaE.** We recombinantly expressed CoaA, CoaD, and CoaE, each with an N-terminal MBP tag, in *E. coli* BL21(DE3), following our previous method[3]. We grew single colonies in LB broth (5 mL) with 50 mg/L kanamycin for 12 h at 37 ºC. We then used cultures to inoculate 1L LB broth with 50 mg/L kanamycin and incubated at 37 ºC until the $OD_{600}$ reached 0.6 to 0.8, before performing cold induction with 0.5 mM IPTG at 18 ºC, followed by shaking for 16 h at 18 ºC and 150 rpm. We obtained cell pellets by centrifugation at 2,000 rpm for 30 min. We stored the cell pellets at -20 ºC or used them directly. We resuspended the cell pellets in cold lysis buffer (50 mM potassium phosphate, pH 7.5, 150 mM NaCl, 10% glycerol) and sonicated on ice for 10 min with 1 s on and 3 s off cycles. We then centrifuged the resulting lysate at 5,000 rpm for 1 h to remove insoluble impurities. We loaded the resulting supernatant onto an amylose column (New England BioLabs), washed it with lysis buffer containing 10 mM maltose, and eluted with lysis buffer containing 100 mM maltose. We pooled the eluted samples and dialyzed them using 10K MWCO SnakeSkin dialysis tubing (Thermo Fisher Scientific) with lysis buffer (1 L) for 12 h at 4 ºC. We concentrated the dialyzed protein to 10 mg/mL and flash-froze it in aliquots for storage at -80 ºC or used it directly.

**Apofication and crosslinker loading of *E. coli* ACP[3]**. We expressed and purified native *E. coli* ACP in a mixture of *apo*- and *holo*- forms. Since Sfp loads substrate on *apo*-ACP only, the conversion of *holo*-ACP into *apo*-ACP was required. We accomplished the conversion using AcpH. To a solution containing 5 mg/mL ACP, 5 mM $MnCl_2$, 10 mM $MgCl_2$, and 0.25% DTT in 50 mM Tris-HCl buffer, pH 7.4, 150 mM NaCl, 10% glycerol, we added 0.01 mg/mL AcpH. We incubated the reaction at 23 ºC for 12 h. We used gel filtration on a HiLoad 16/600 Superdex 75 pg column (GE Biosciences) to obtain pure *apo*-ACP from the mixture. The purified sample was concentrated to 5 mg/mL using a 3K Amicon Ultra spin concentrator (Millipore). We aliquoted, flash-froze, and stored the resulting sample at -80 ºC, or used it directly in the next crosslinker loading step. For crosslinker loading onto *apo*-ACP, we used a previously developed protocol (Fig. S5). Briefly, the reaction included 1 mg/mL *apo*-ACP, 0.01 mg/mL CoaA, 0.01 mg/mL CoaD, 0.01 mg/mL CoaE, 0.04 mg/mL Sfp, 12.5 mM $MgCl_2$, 1 mM DTT, 8 mM ATP, and 0.2 mM crosslinker (50 mM DMSO stock solution) in 50 mM Tris-HCl, pH 7.4, 150 mM NaCl, 10% glycerol. We controlled the final DMSO concentration to be below 1%. We allowed the reaction to incubate for 12 h at room temperature, followed by gel filtration through a HiLoad 16/600 Superdex 75 pg (GE Biosciences) size exclusion column using FPLC (ÄKTA Pure) with 50 mM Tris-HCl, pH 7.4, 150 mM NaCl, 10% glycerol to yield pure crosslinker-loaded ACP or *crypto*-ACP. We concentrated the *crypto*-ACP to 5 mg/mL using a 3K spin concentrator (Millipore Amicon Ultra). We aliquoted, flash-froze, and stored the resulting sample at -80 ºC.

**Crosslinking between *crypto*-ACP and *E. coli* ketosynthases**. We incubated *crypto*-ACP loaded with crosslinker **1-8** (150 μM) with FabF, FabFI108F, FabFG198F, FabB, and FabBG107M (30 μM) in a buffer containing 50 mM Tris-HCl, pH 7.4, 150 mM NaCl, 10% glycerol at 25 ºC for 16 h. We evaluated the crosslinking reactions using SDS-PAGE analyses.

**X-ray crystallography.** Prior to crystallization, we submitted both $FabB_{G107M}$=$ACP_{C8}$ and $FabB_{G107M}$=$ACP_{C12}$ to Hiload Superdex 200 pg (16/600) size exclusion column using FPLC (ÄKTA Pure) with 50 mM Tris-HCl, pH 7.4, 150 mM NaCl, 10% glycerol to remove aggregated protein. We then buffer-exchanged the purified crosslinked complexes into minimal buffer containing 20 mM Tris, pH 7.4, 50 mM NaCl using a 30K spin concentrator (Millipore Amicon Ultra) to 9 mg/mL. Briefly, for the crystallization of $FabB_{G107M}$=$ACP_{C8}$ and $FabB_{G107M}$=$ACP_{C12}$, we added 1 μL of crosslinked complex to 1 μL of crystallization solution (0.1 M sodium cacodylate, pH 6.0, 0.3 M sodium acetate, 28% PEG 8000), then incubated at 4 ºC using a hanging-drop vapor-diffusion method. We incubated the crystals in LV CryoOil (MiTeGen) briefly before freezing in liquid nitrogen. We collected data for both crosslinked complexes through Advanced Light Source beamline 5.0.3 at 100 K and 0.999 Å. We indexed, scaled, and merged diffraction data through iMosflm[4] and Aimless[5] within the CCP4i2 software suite[6]. We performed molecular replacement through PHASER[7] using wtFabB=$ACP_{C12}$ (PDB: 6OKC). We performed all modeling, map refinement, and covalent crosslinker linking using Coot, REFMAC5[8], and AceDRG[9]. We deposited the processed density maps and structures on the Protein Data Bank under entries PDB: 9MLW for $FabB_{G107M}$=$ACP_{C8}$ and PDB: 9MLX for $FabB_{G107M}$=$ACP_{C12}$, with the statistics shown in Table S1.

***In vitro* analysis of fatty acid synthases.** We characterized variants of FabF and FabB by swapping them into the *in vitro* reconstituted FAS from E. coli. To focus on the contribution of FabF, we used a FabB-less reference system (1 μM of a FabF variant, 0 μM FabB, 1 μM all other Fab enzymes, 10 μM TesA, 10 μM holo-ACP, 10 mM HEPES, pH 7.4, 150 mM NaCl, 0.5 mM TCEP) supplemented with 2.6 mM NADPH or NADH, 1.5 mM malonyl-CoA, and 300 μM acetyl-CoA. We initiated reactions by mixing proteins, substrates, and cofactors, then let reactions sit at room temperature (~21 ºC) for 2 hours. We prepared extracted fatty acids for GC/MS analysis using the procedure described in prior work[1]. We quantified fatty acids present in unknown samples with standard curves for each fatty acid. For each composition, we used at least three independently reconstituted reactions.

**Molecular dynamics.** For all MD simulations, we derived the FabB-acyl complexes from the crystal structure of FabB covalently bound to a C12 acyl chain (PDB 1EK4), the FabB-ACP complex from the crystal structure of the cross-linked FabB–acyl-ACP complex with a C16 acyl chain (PDB 6OKF). The FabF-acyl and ACP complexes were derived from the Apo FabF dimer structure (PDB 2GFW). We used MODELLER to introduce the G107M mutation and Avogadro to shorten or elongate the acyl chain to generate C4-16 initial configurations. The homology between FabF and FabB was used to align the acyl-intermediate and ACP complex structures in FabB to the apo FabF structure to get initial configurations. All systems were parameterized using the Amber ff14SB protein force field and GAFF2 parameters for the acyl chain. The unit cell boxes for all simulations conducted in solvent were constructed to maintain a minimum distance of 1 nm between the molecule and the periodic boundary and then filled with TIP3P solvent. All simulations were run using GROMACS 2025.0. The equilibration procedure was identical for all systems. This procedure consisted of an initial energy minimization to a threshold of 50 kJ/mol*nm followed by a 100 ps NVT equilibration using the Bussi-Parrinello thermostat maintaining a temperature of 300 K. A 100 ps NPT equilibration was also performed with the stochastic cell rescaling barostat for all calculations performed in solvent to maintain a pressure of 1 atm. All simulations were performed with a 2-fs time step. The procedure for extracting decorrelated initial configurations for each replica is detailed in our previous work[10].

The process for segmenting trajectory frames into pockets A, B, and C is detailed in our previous work for the covalent intermediate. For the ACP complex, we instead used acyl chain RMSD to align reference frames selected by manual pocket identification. First, we manually identified 25 frames in pocket A, B, and C from both the FabB and FabF ACP complexes for chain lengths 4, 6, 8, 10, 12, 14, and 16. For each pocket, we computed the centroid of each set as the reference structure, then computed the RMSD for each reference frame relative to that pocket's reference structure. A frame was labeled as being in a designated pocket if its RMSD to that pocket's reference structure was the lowest among the three and also below a set threshold. We determined the thresholds by computing the maximum RMSD for manually identified pocket A, B, or C structures (Figure S8). These thresholds are 0.85, 0.8, and 0.8 for pockets A, B, and C, respectively. We computed the alchemical free energy (FE) for each transformation using the procedure outlined in our previous work. In short, the apo acyl-ACP configuration was used as the reference state for all FE transformations, and the simulations were segmented by trajectory frame into pockets A, B, and C before the FE was computed using MBAR. We also performed additional trajectory analysis to compute non-bonded and h-bond interactions. Non-bonded interactions were defined as frames in which the heavy atom distance between two atoms was less than 5 Å, and h-bonds were defined by a H-acceptor distance of less than 2.5 Å and a donor-H-acceptor angle greater than 120°.

**SI FIGURES**

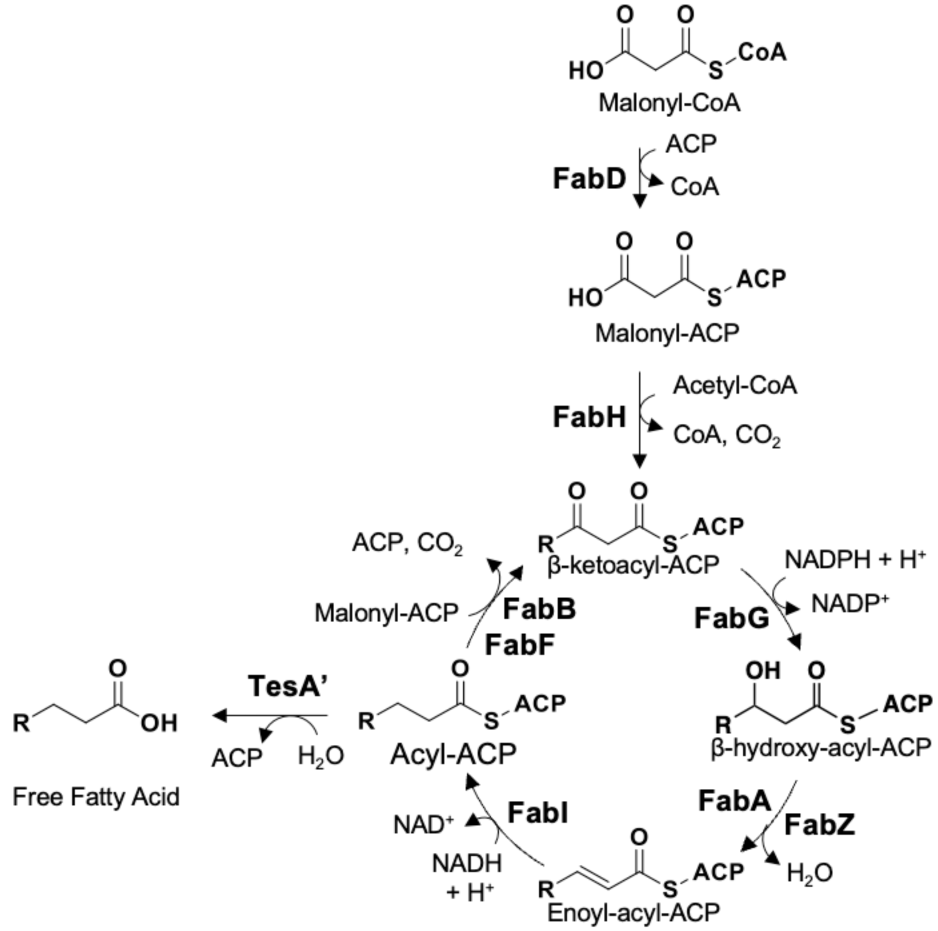


**Figure S1. Fatty acid synthesis in *E. coli.*** A depiction of the fatty acid synthase (FAS) of *E. coli* supplemented with a leaderless variant of TesA (TesA').

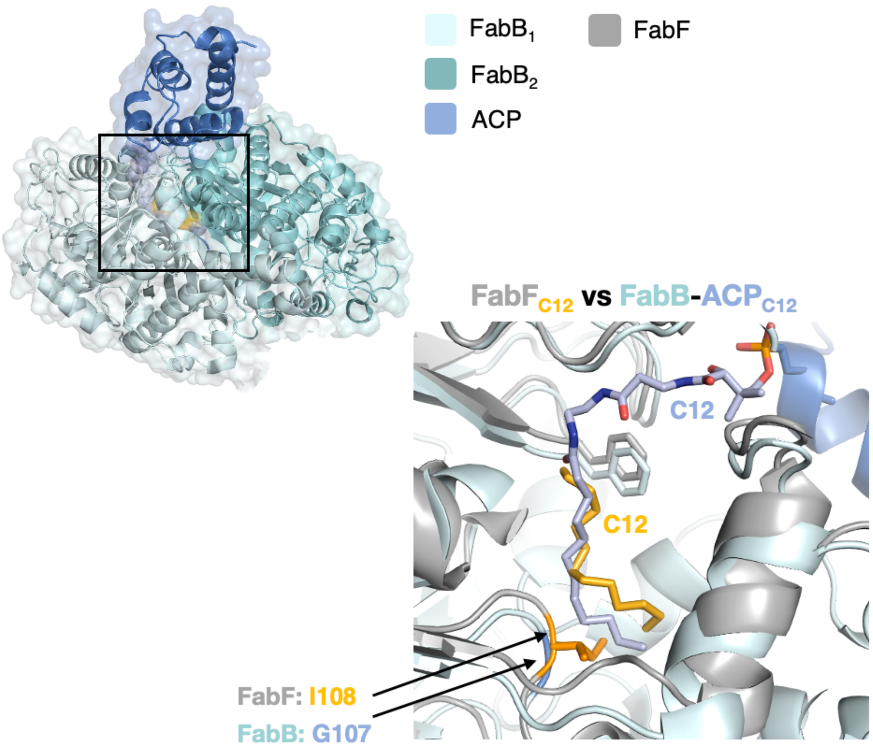


**Figure S2. Comparison of FabB and FabF.** Overlay of X-ray crystal structures of $FabF_{K335A}$ with covalently linked dodecanoic acid (PDB entry 2GFY) and FabB with covalently crosslinked $ACP_{C12}$ (N-[2-(dodecanoylamino)ethyl]-N~3~-[(2R)-2-hydroxy-3,3-dimethyl-4-(phosphonooxy)butanoyl]-beta-alaninamide; PDB entry 6OKF). This overlay does not explain why FabB is more permissive of G107M than FabB is of I107M.

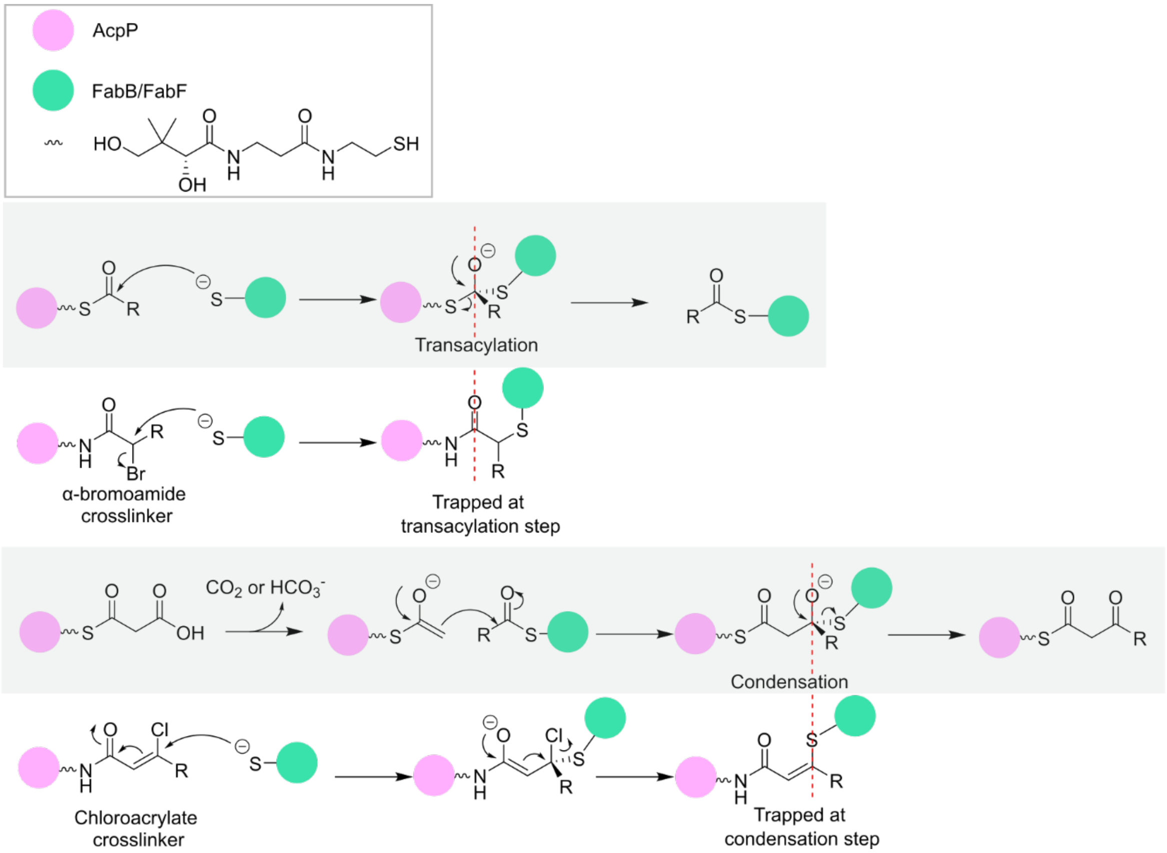


**Figure S3. Trapping distinct catalytic states in FabB/FabF.** KSs catalyze acyl chain elongation in two steps: transacylation and condensation (grey shading). Both steps form a tetrahedral intermediate. In prior work[11], we developed natural substrate-mimicking crosslinkers that covalently crosslink ACP and KS, thereby trapping the transacylation and condensation steps and yielding products that resemble the tetrahedral intermediates.

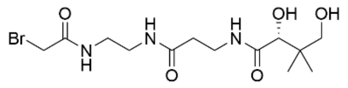

**1**

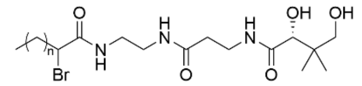

**2** n=1
**3** n=3
**4** n=5
**5** n=9
**6** n=13

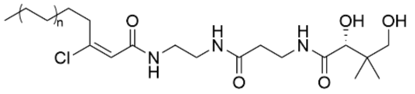

**7** n=1
**8** n=4

**Figure S4. Chemical structure of crosslinkers.** A panel of crosslinkers equipped with an α-bromo motif (**1-6**) or a chloroacrylate motif (**7-8**), both of which target the catalytic cysteines of FabB and FabF.

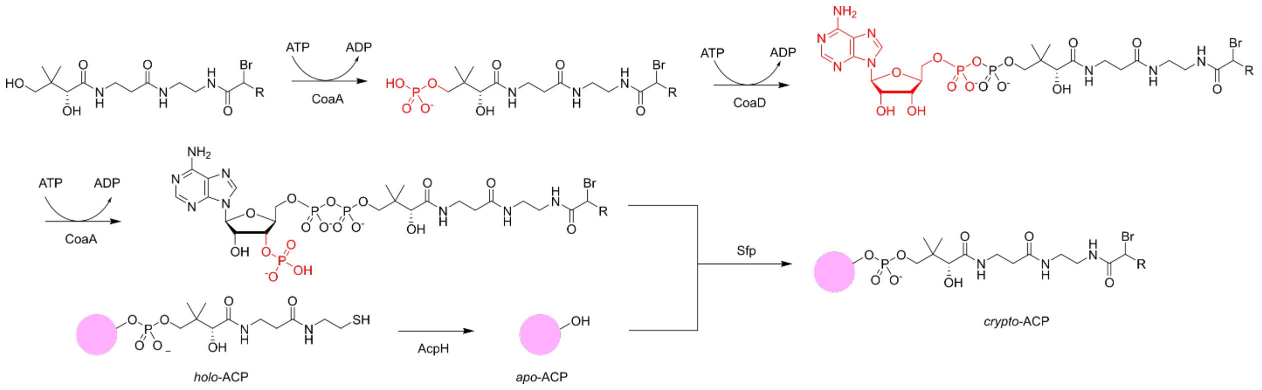


**Figure S5. Crosslinker loading onto *apo*-ACP using a one-pot enzymatic reaction.** This one-pot enzymatic reaction enables crosslinker loading onto *apo*-ACP. Briefly, crosslinkers are phosphorylated, adenylated, and phosphorylated sequentially by enzymes CoaA, CoaD, and CoaE, respectively, to achieve primed crosslinkers. AcpH converts *holo*-ACP to *apo*-ACP (pink dot is ACP), and Sfp recognizes and loads primed crosslinkers onto the active residue Ser of *apo*-ACP to yield crosslinker-loaded ACP, hereafter *crypto*-ACP.

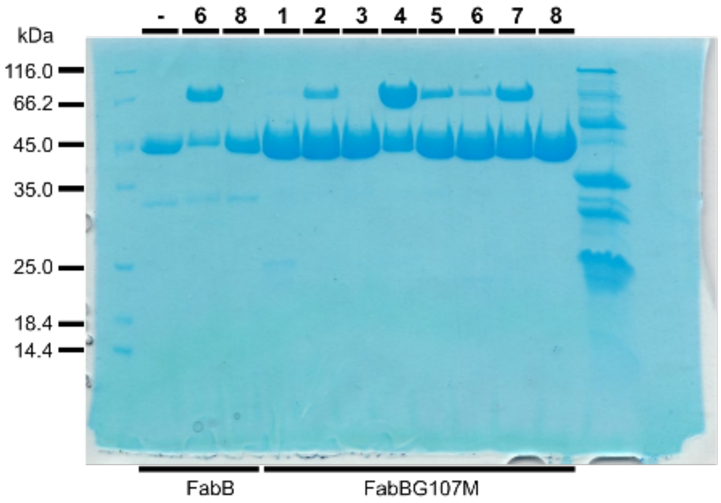


**Figure S6. SDS-PAGE analysis of crosslinked FabB-ACP.** SDS-PAGE (12%) analysis of $FabB_{WT}$ or $FabBG_{107M}$ crosslinked with ACP. A brief summary of crosslinkers (with acyl chain lengths): α-bromopantetheine amide crosslinkers 1 (C2), 2 (C4), 3 (C6), 4 (C8), 5 (C12), and 6 (C16); and chloroacrylate crosslinkers 7 (C8) and 8 (C14). See Fig. S4 for structures. Bands show FabB (45.0 kDa) or FabB-ACP complexes (66.2 kDa).

a) b) c) d) e) f) g) h)

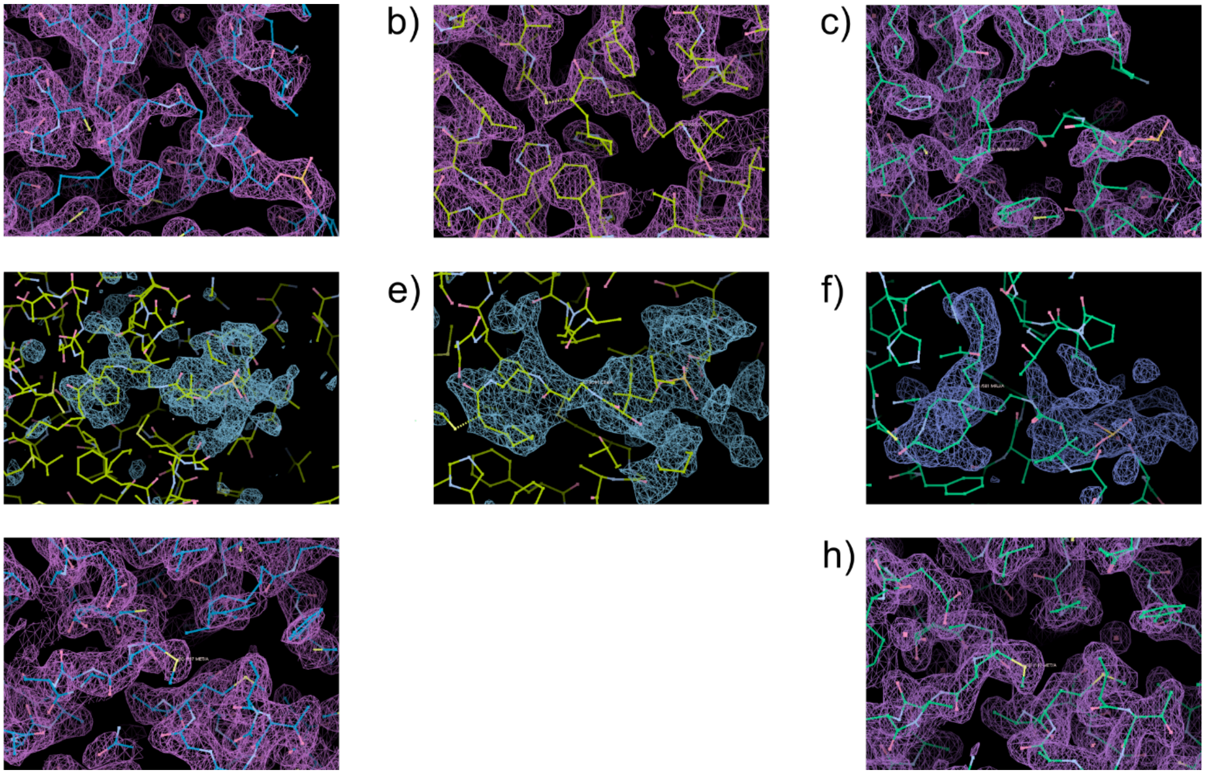

**Figure S7. Structural analysis of $\mathbf{FabB_{G107M}{=}ACP_{C8}}$ and $\mathbf{FabB_{G107M}{=}ACP_{C12}}$.** (a-b) Density maps of $FabB_{G107M}{=}ACP_{C8}$ ($\sigma = 1$) allow unambiguous positioning of the pantetheine arm of crosslinker **4** but not the acyl tail. (c) Density maps of $FabB_{G107M}{=}ACP_{C12}$ ($\sigma = 1$) support the acyl tail of crosslinker **5**, but the pantetheine arm is challenging to resolve. (d-e) Polder maps for $FabB_{G107M}{=}ACP_{C8}$ ($\sigma = 2.5$) suggest that the acyl tail adopts two conformations, which are consistent with pockets A and B. (f) The polder map ($\sigma = 3$) of $FabB_{G107M}{=}ACP_{C12}$ shows an acyl chain in pocket B. (g-h) The map densities support the G107M mutation and truncation of pocket A for both structures.

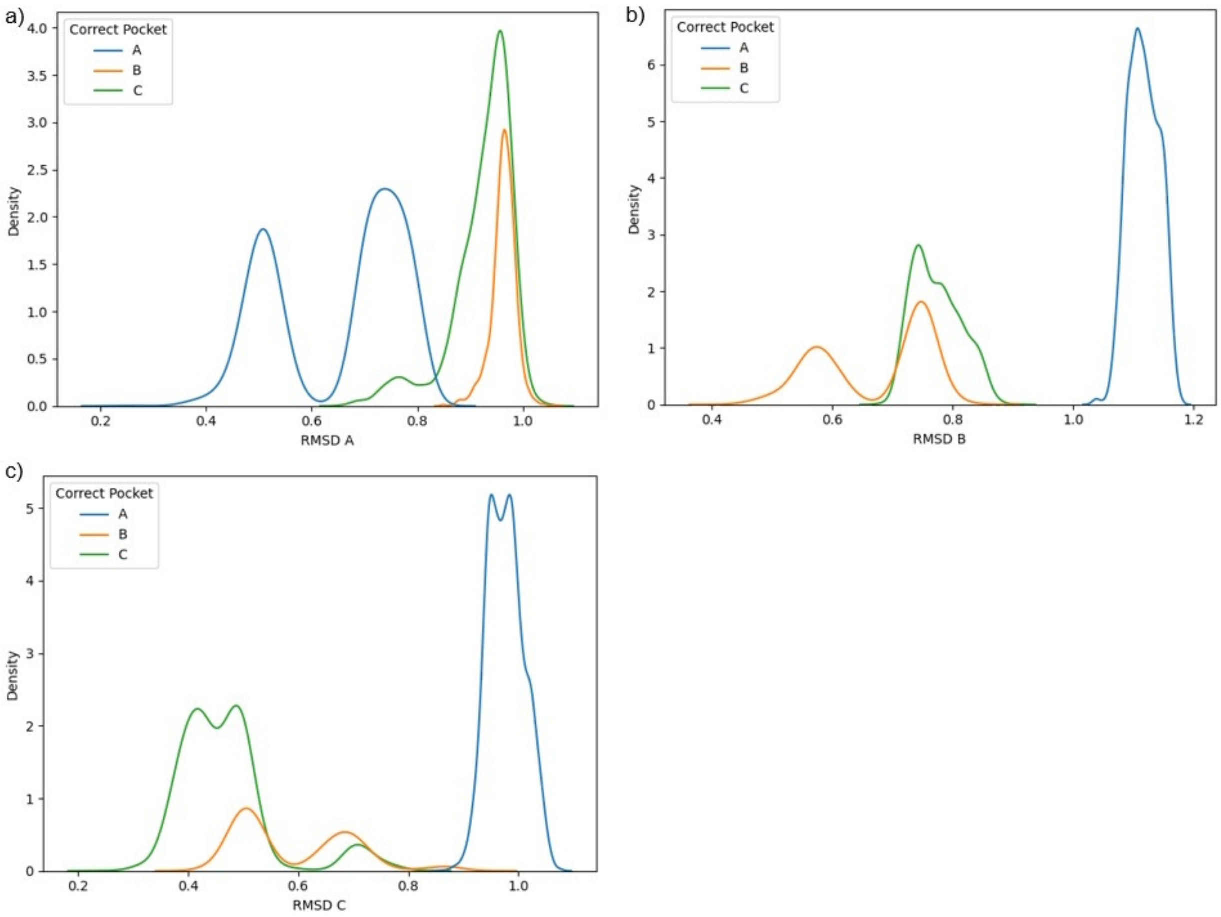


**Figure S8. RMSD to pocket references.** Plots of the RMSD between manually selected frames with the acyl chain in pockets A, B, and C, relative to reference structures for pockets (a) A, (b) B, and (c) C. For each figure, we selected 25 frames with the acyl chain in the designated pocket of FabB-ACP and FabF-ACP for all chain lengths of 4, 6, 8, 10, 12, 14, and 16 (i.e., 25 frames per chain length per complex). We extracted the frames from 100 ns MT-REXEE simulations of the first replicate of each system and calculated the RMSD of heavy atoms in the acyl chain. These results allowed us to establish RMSD thresholds to automate pocket sorting by trajectory frame.

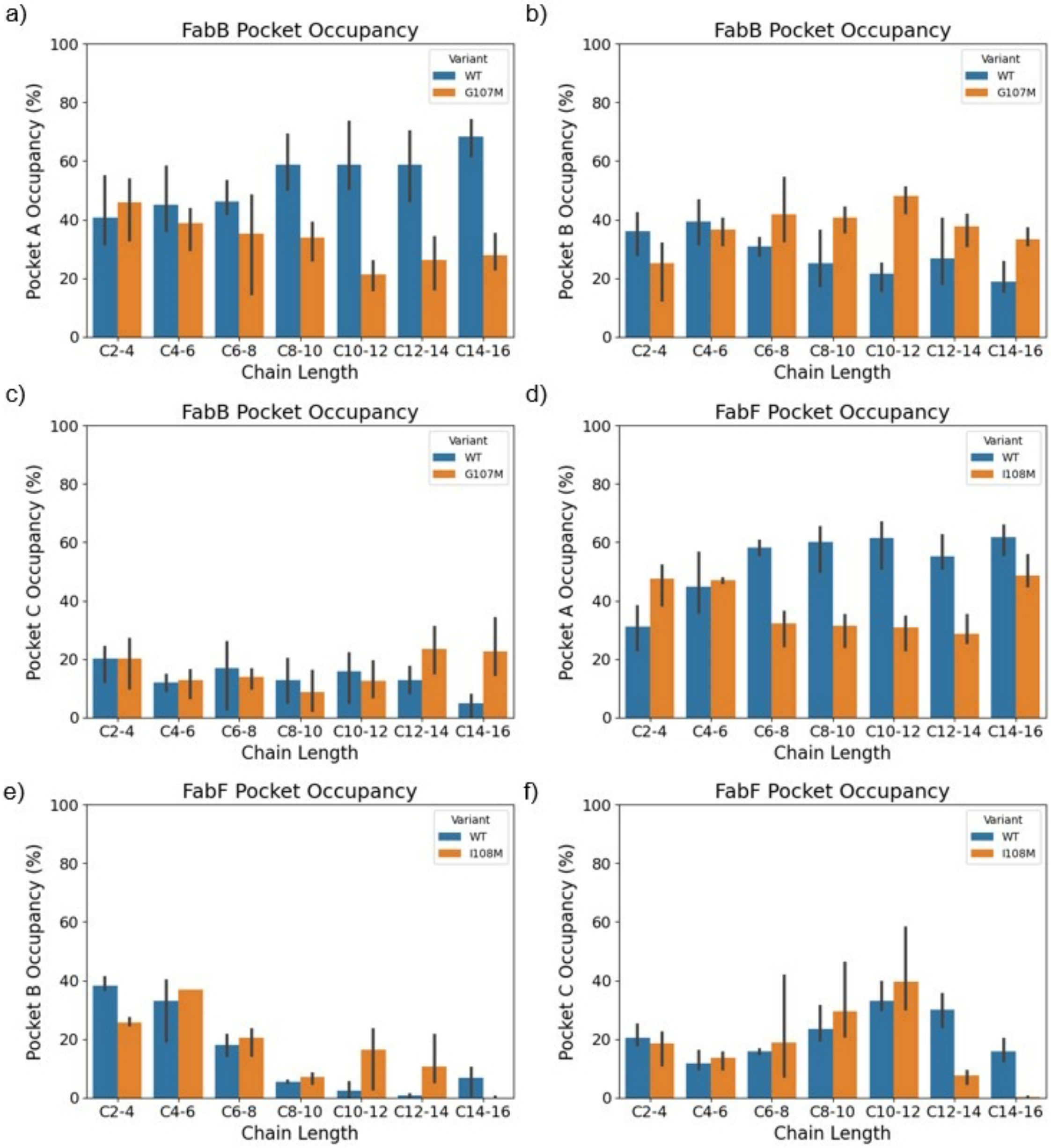


**Figure S9. Pocket occupancy in MT-REXEE simulations of FabF-ACP and FabB-ACP complexes.** (a)WT FabB uses Pocket A, the canonical binding pocket, as the primary pocket for all chain lengths. The G107M mutation blocks pocket A for substrates with 8 or more carbons, decreasing the occupancy of pocket A and (b) increasing the occupancy of pocket B for these longer chains. (c) The G107M mutation also increases the occupancy of pocket C for long chains, though pocket B remains the dominant pocket for all acyl chains longer than eight carbons. (d) In FabF, the I108M mutation causes a similar decrease in the occupancy of pocket A, but (e) does not enhance sampling of pocket B or (f) pocket C. Instead, we observe a shift in ACP that results in a misalignment of the catalytic carbonyl on the acyl chain and CYS163, yielding a non-catalytically competent conformation. Data represent the mean and standard error of n = 3 simulations of each chain length.

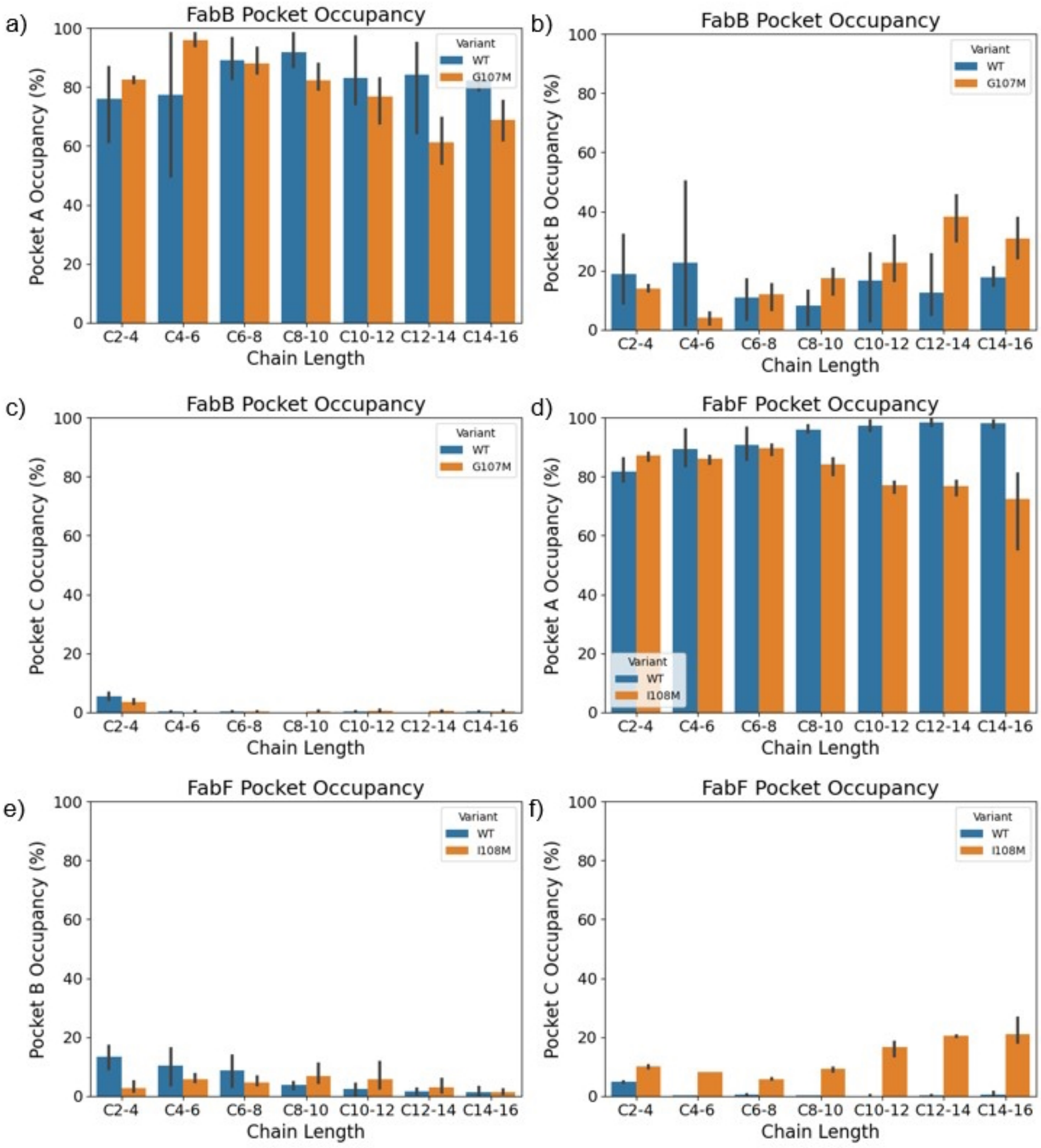


**Figure S10. pocket occupancy in FabF and FabB acyl intermediate complex MT-REXEE simulations.** (a)In FabB, the G107M mutation still shifts binding away from pocket A in (b) favor of pocket B with (c) minimal sampling of pocket C. (d) In FabF, the analogous mutation decreases occupancy of pocket A and (e) increases sampling of pocket B very slightly with (f) long chains binding preferentially to pocket C, and less time in a catalytically competent conformation overall. Data represent the mean and standard error of n = 3 simulations of each chain length.

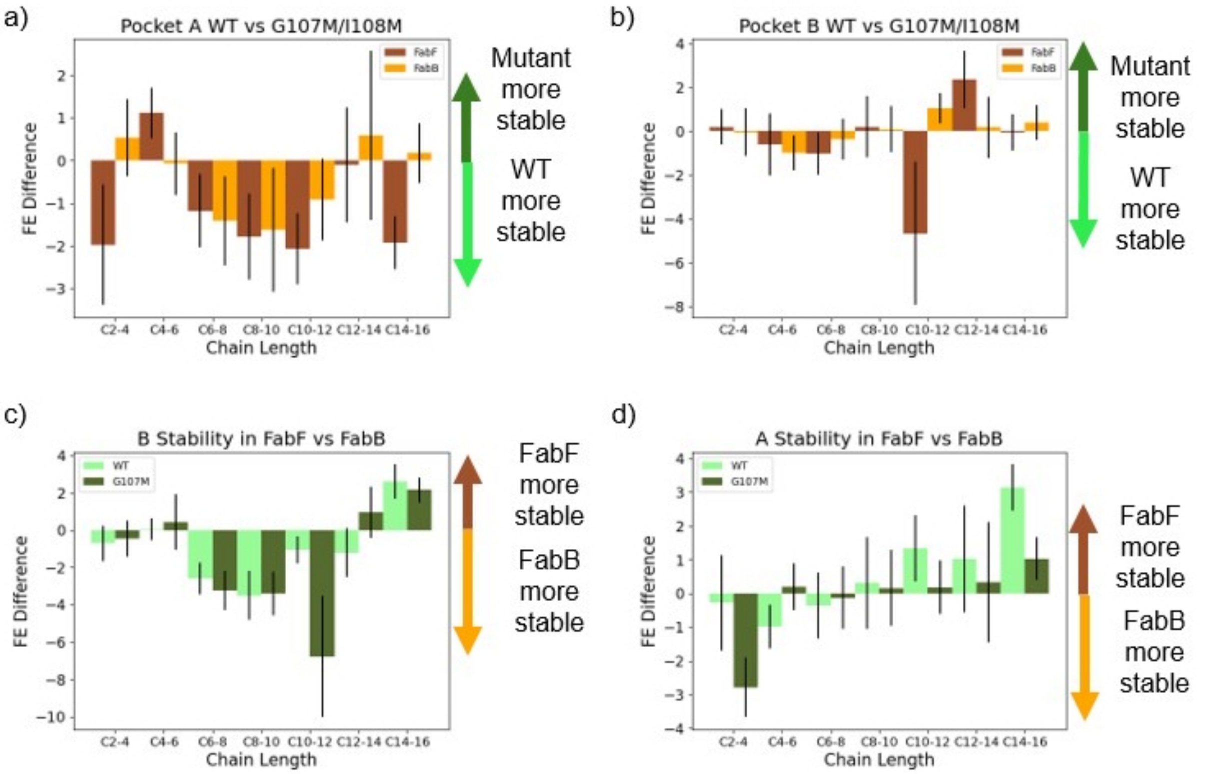


**Figure S11. Alchemical binding FE estimates in FabF and FabB ACP complex.** These plots depict the alchemical relative binding FE difference between the free acyl-ACP and the acyl-ACP in complex with FabB or FabF for incremental addition of two acyl chain carbons. (a) Mutating residues FabB 107 or FabF 108 to Met yields a 2-3 kcal/mol penalty for the extension of the acyl chain into pocket A for the C6-8, C8-10, and C10-C12 transformations. (b) The relative stability of pocket B is within statistical error and statistically equivalent ($p > 0.05$) between WT and G107M for all transformations for both FabB and FabF, except for C10-12 and C12-14 in FabF. This suggests that there is little to no allosteric effect from the mutation, which would increase the stability of pocket B. (c) Pocket B is, however, 3-7 kcal/mol more stable in FabB than in FabF. The significant difference for the C6-8 transformation indicates that insertion of an acyl chain of 8 carbons or longer into pocket B in FabF is unlikely to be stable. (d) Pocket A stability is similar for FabF and FabB regardless of variant. Pocket A is moderately more stable in FabF WT compared to FabB WT for chain lengths greater than 8 carbons. The significance of this observation is uncertain, and given that the same effect does not appear between the FabF and FabB mutant variants and there is no evidence for an allosteric effect from the mutation, this observation is probably not significant. Data represent the mean and standard error of $n = 3$ simulations of each chain length.

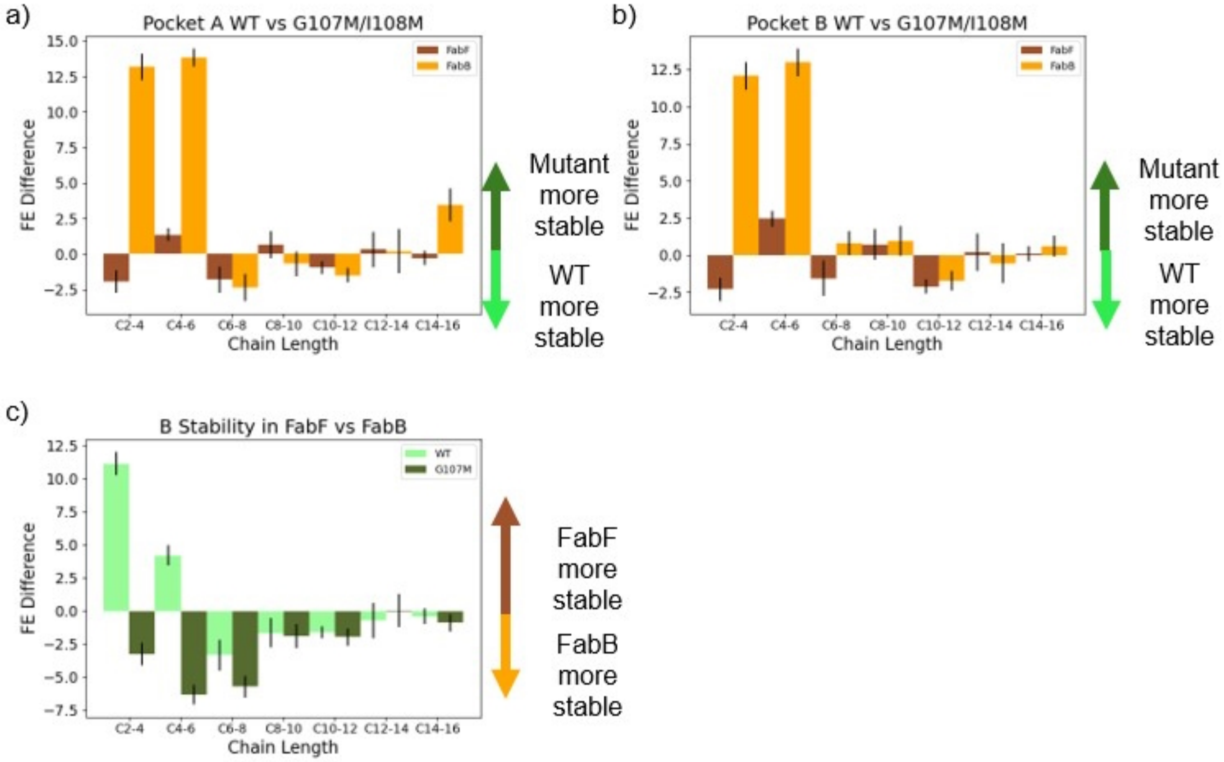


**Figure S12. Alchemical binding FE estimates in FabF and FabB acyl intermediate complex.** These plots depict the alchemical relative FE difference between free acyl-ACP and acyl-FabF or acyl-FabB (covalently attached acyl chain) as 2 carbons are incrementally added to the acyl chain. (a) Mutating residues FabB 107 or FabF 108 to Met yields a 1.5-2 kcal/mol penalty for the extension of the acyl chain into pocket A for the C6-8 transformation. (b) The relative stability of pocket B is within statistical error and statistically equivalent ($p > 0.05$) between WT and G107M for all transformations for both FabB and FabF, except for C10-12 for both FabB and FabF. Only one transformation exhibits a moderately significant difference ($0.01<p<0.05$), which is insufficient for allostery. (c) Pocket B is 3-5 kcal/mol more stable in FabB than FabF. The significant difference for the C6-8 transformation indicates that insertion of an acyl chain of 8 carbons or longer into pocket B in FabF is not stable. Data represent the mean and standard error of n = 3 simulations of each chain length.

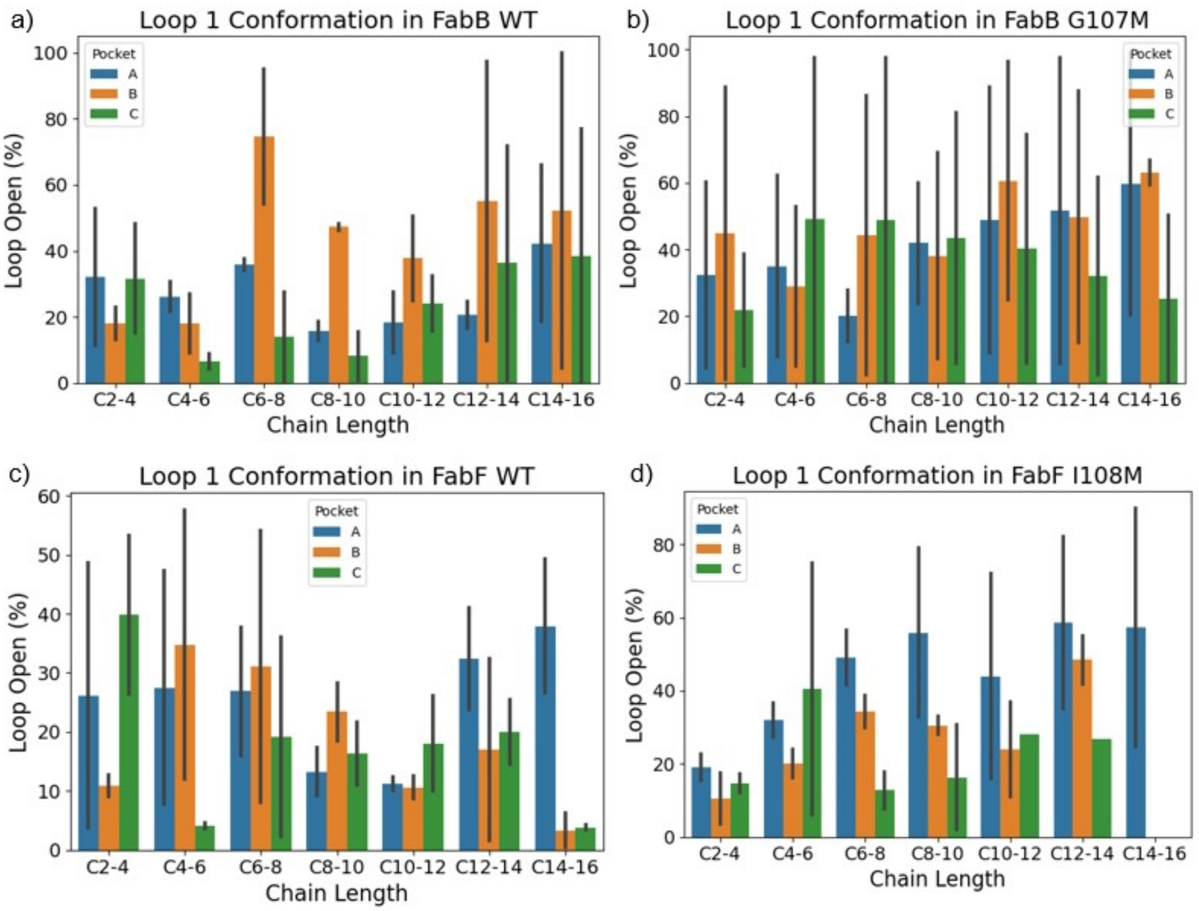


**Figure S13. Evaluation of gate open conformation in ACP complex.** All simulations were initiated with the gating loop 1 in the closed conformation, but throughout the simulation, the loop is free to reorient. This loop is highly flexible and reorients between open and closed relatively frequently throughout the simulations. There is no correlation between loop opening and the acyl chain being present in either pockets A, B, or C. In general, the loop is open more in the mutant variants, though these differences are not statistically significant ($p > 0.05$). Data represent the mean and standard error of $n = 3$ simulations of each chain length.

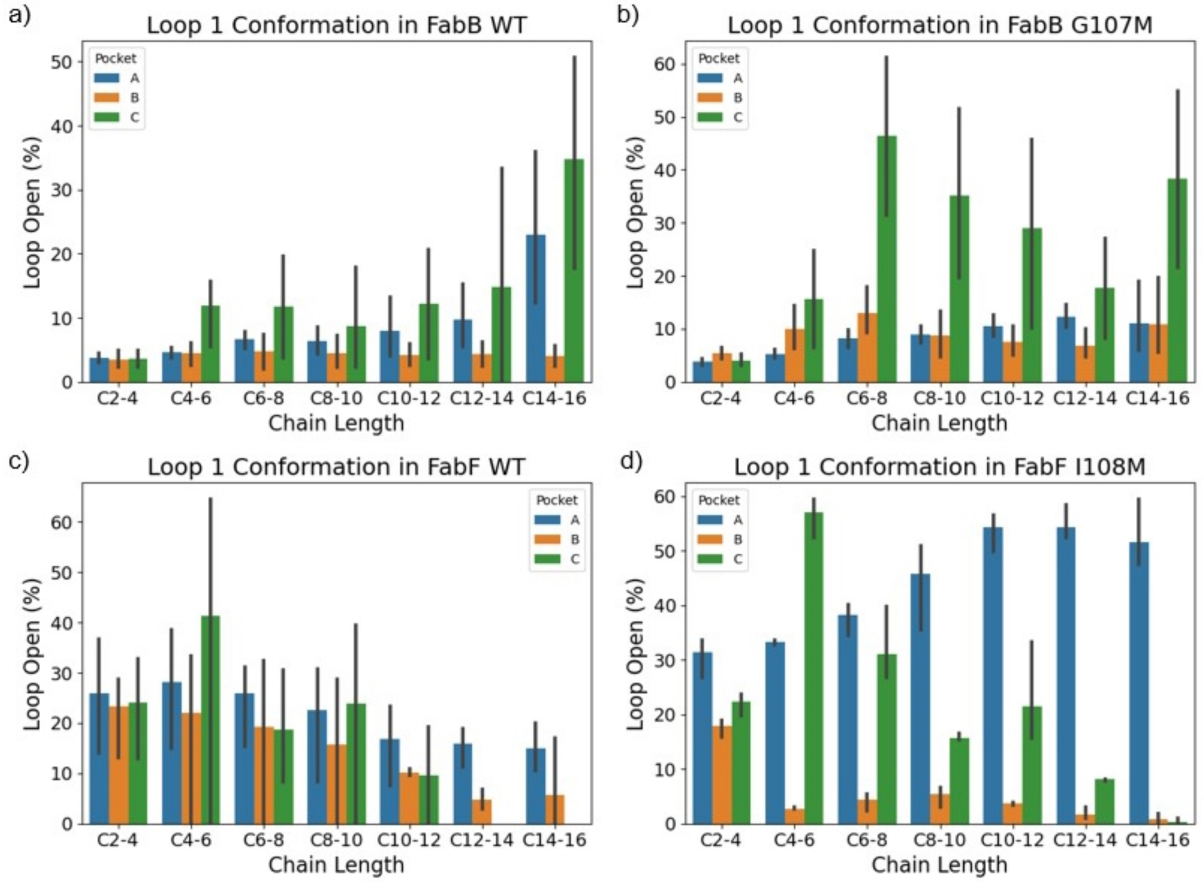


**Figure S14: Evaluation of gate open conformation in the covalent intermediate.** All simulations were initiated with gating loop 1 in the closed conformation, but throughout the simulation, the loop is free to reorient. The loop is flexible throughout simulations, particularly for longer-chain substrates. The loop also reorients to the open conformation significantly more in FabF than in FabB. Given the high degree of variation between replicas and the fact that no simulation reorients to the open conformation for a majority of the trajectory, this effect seems minor. Our main conclusion is that the observed differences in pocket occupation and binding FE are an average of the open and closed conformation and do not correspond exclusively to one conformation. Data represent the mean and standard error of n = 3 simulations of each chain length.

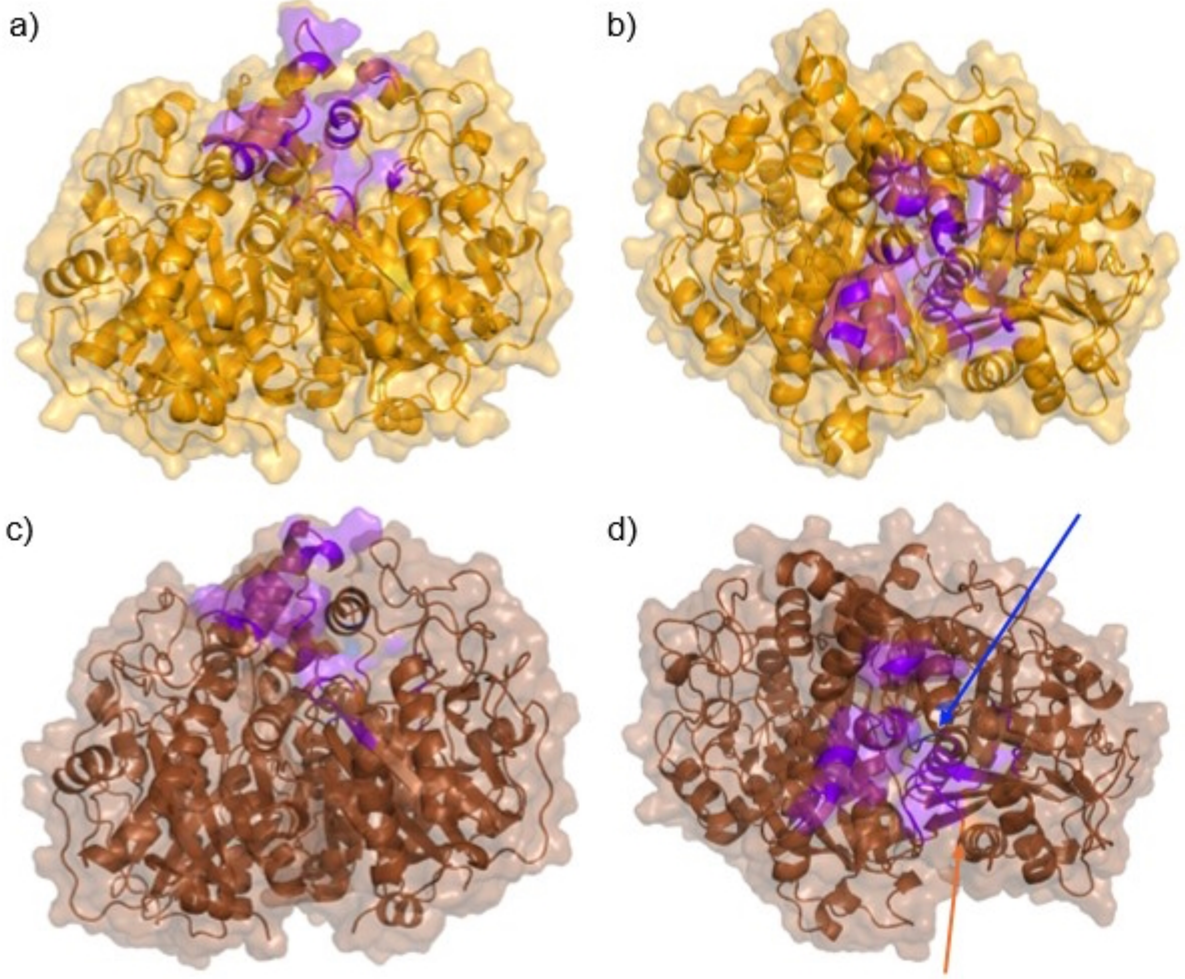


**Figure S15. Comparison of protein-protein interactions at the Fab-ACP Interface.** This figure shows the residues in (a-b) FabB and (c-d) FabF that form non-bonded interactions with ACP for a minimum of 60% of MD trajectories for all substrate chain lengths in both the WT and G107M/I108M variants. Colored residues show interactions formed primarily when the acyl chain is in (orange) pocket A, (blue) pocket B, or (purple) both. There were only two pocket-dependent differences in the ACP interface (highlighted by arrows in their respective colors), and as such, both interfaces are primarily purple. The Fab-ACP interface shows no major differences when the acyl chain occupies pockets A or B.

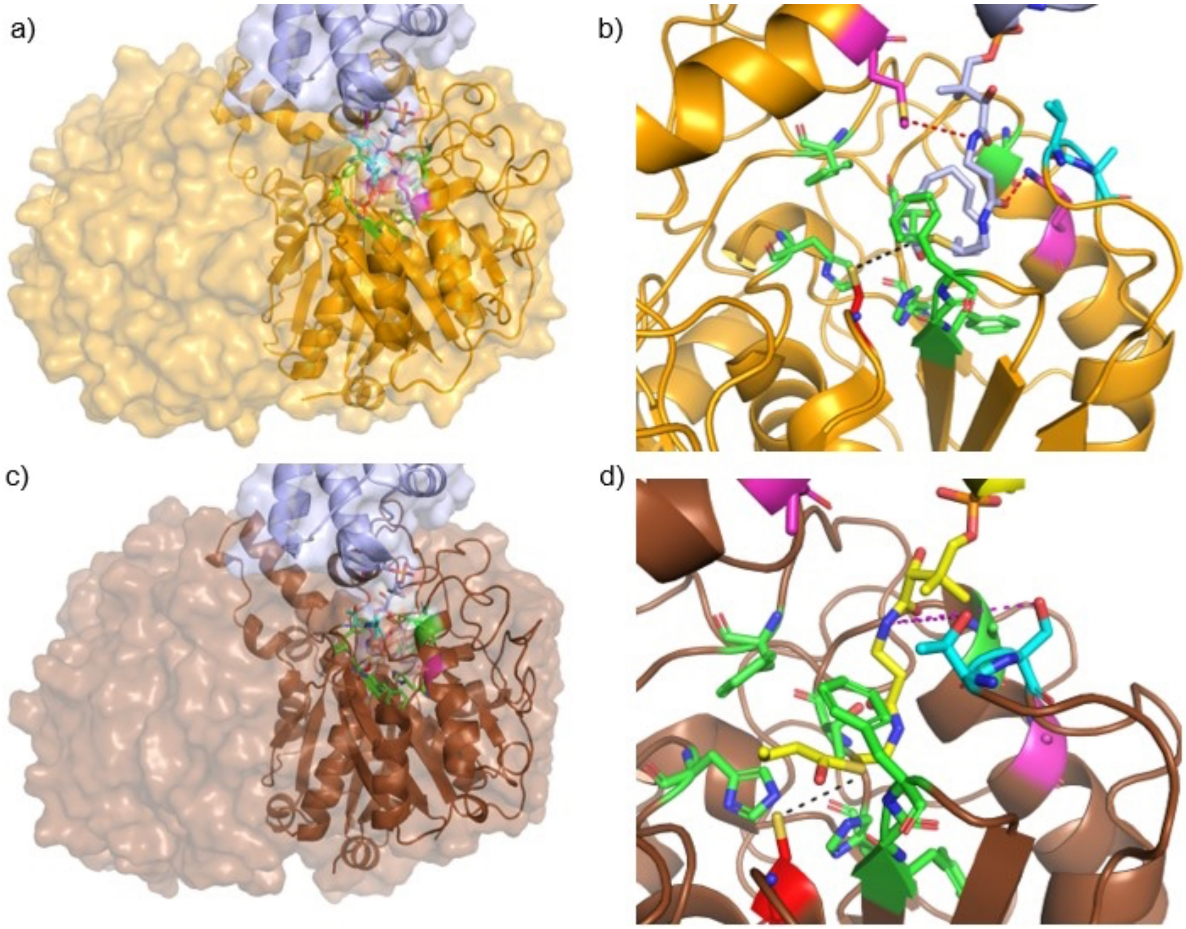


**Figure S16. Comparison of KS-Ppant interactions.** These figures highlight the interactions formed between KS and the Ppant linker when the acyl chain is in pocket B for (a-b) FabB and (c-d) FabF. The catalytic Cys is shown in red with a black dotted line connecting it to the catalytic carbonyl on the acyl chain. Residues in green are identical or highly similar between analogous positions in FabB and FabF. Residues Met206 and Lys310 (pink) engage in h-bonds with the Ppant linker in FabB; their FabF counterparts, Ala204 and A312, are unable to form such interactions. H-bonds that form for >75% of trajectories appear as red dotted lines; h-bonds that form >50% of trajectories, pink dotted lines. In FabB, Val272 and Ala273 (blue) form weak transient interactions with the Ppant linker when the acyl chain occupies pocket B, while in FabF, the corresponding residues Thr270 and Ser271 form h-bonds that may limit linker flexibility.

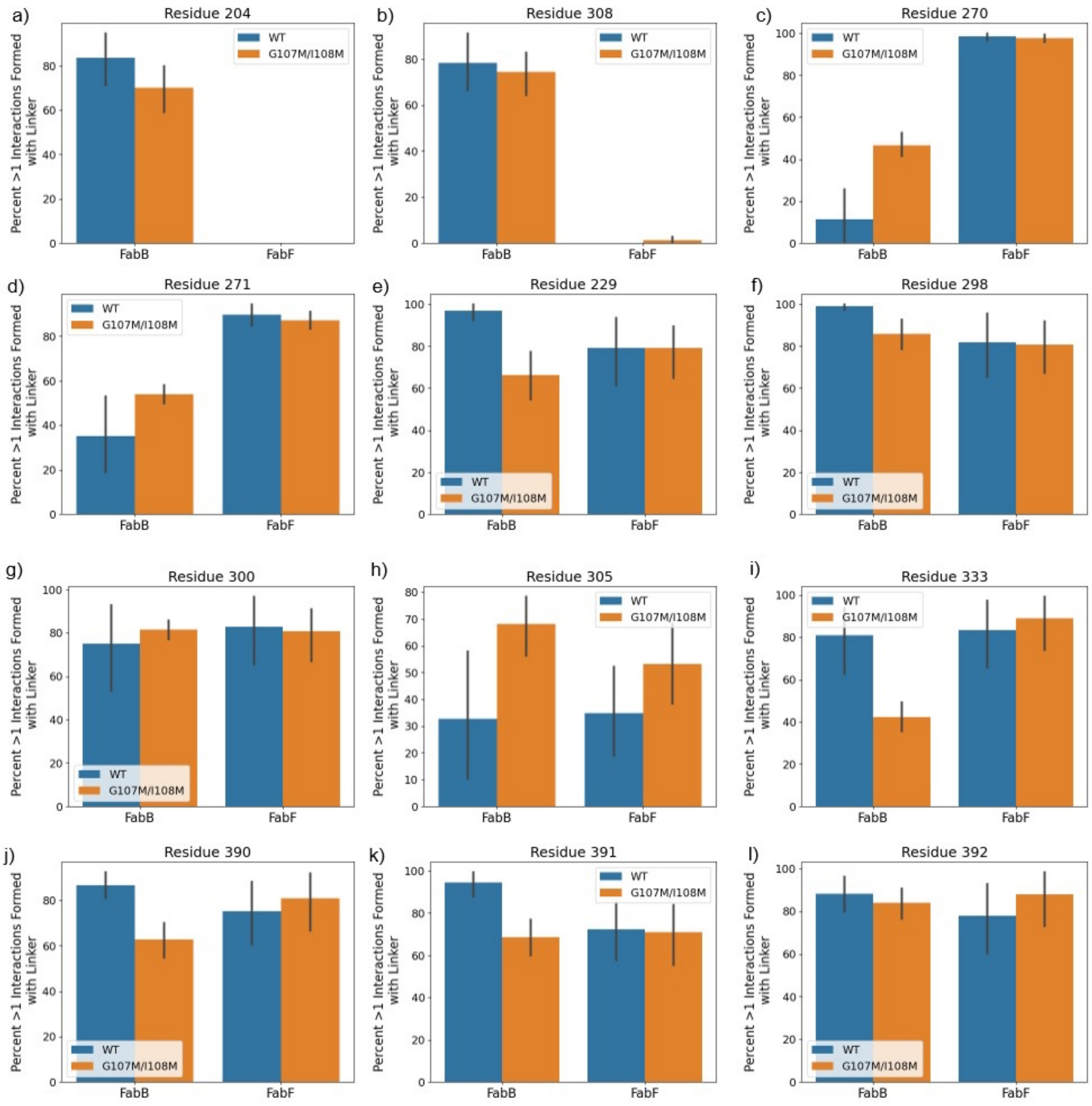


**Figure S17. Comparison of KS interactions formed with the Ppant linker.** We compare the percentage of each trajectory for which any non-bonded interaction is formed between a given FabF/FabB residue and the Ppant linker when the acyl chain occupies pocket B. The mean averages three independent replicas, and the error bars represent the standard error of the mean. The residue numbers correspond to FabB and the analogous position on FabF, as determined by structural alignment. (a-b) Residues Met206 and Lys310 in FabB form consistent h-bonds with the Ppant linker, while equivalent residues Ala204 and Ala312 in FabF do not. (c-d) Residues Val272 and Ala273 in FabB form unstable VDW interactions with the Ppnt linker, while equivalent residues Thr269 and Ser270 of FabF form stable h-bond interactions with this linker. (e-l) The remaining residues in FabB(FabF)—Phe231(Phe228), His299(His302), Thr301(Thr304), Gly306(Ala308), His335(His339), Phe392(Phe397), Gly393(Gly298), and Phe394(His399)—form relatively stable non-bonded interactions with the linker in both enzymes. Data represent the mean and standard error of n = 3 simulations of each chain length.

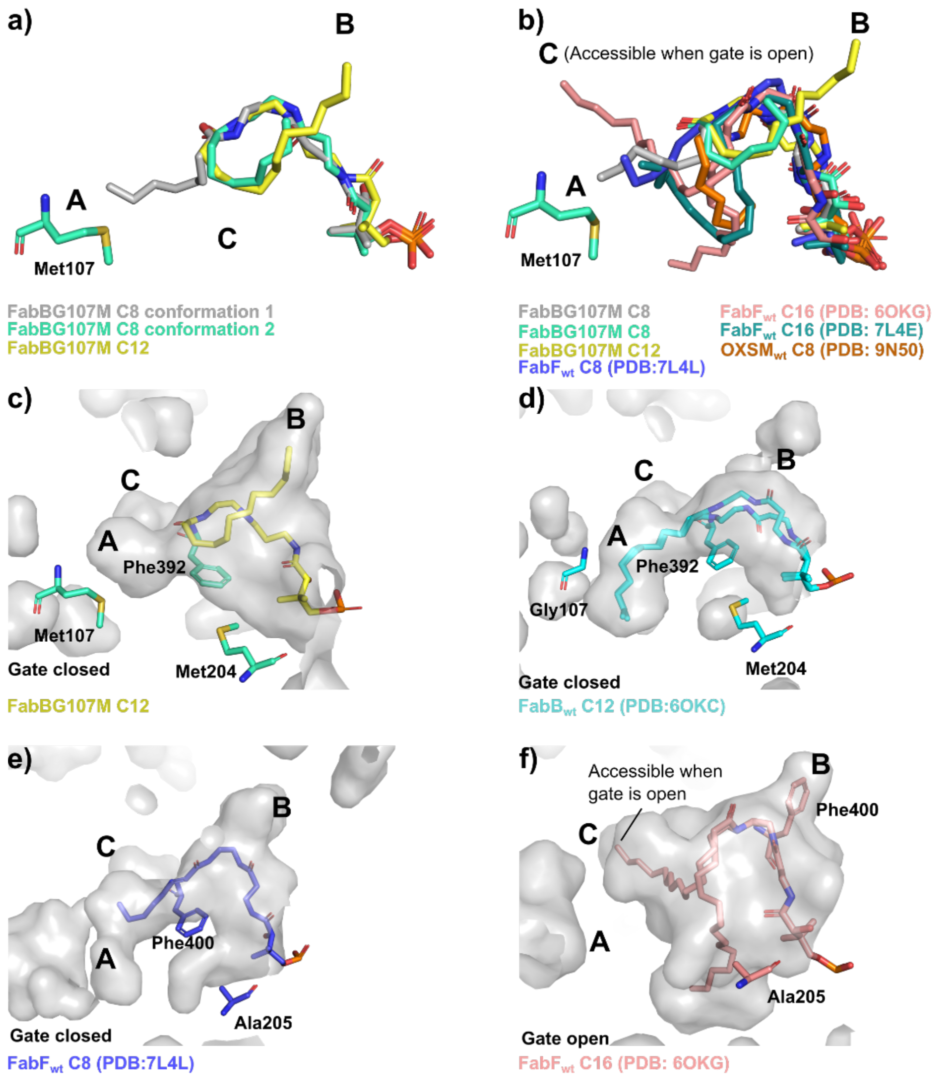


**Figure S18.** Analysis of substrate poses and binding pockets in FabB, FabB$_{G107M}$, and FabF. (a) An overlay of FabB$_{G107M}$=ACP$_{C8}$ with FabB$_{G107M}$=ACP$_{C12}$ (RMSD= 0.237) demonstrates that in FabB$_{G107M}$=ACP$_{C8}$, crosslinker **4** adopts conformation 1 (grey) and conformation 2 (light green). In conformation 1, **4** occupies pocket A, the canonical binding pocket, which is truncated by the G107M mutation. In conformation 2, **4** points toward the binding pocket B. In FabB$_{G107M}$=ACP$_{C12}$, crosslinker **5** (yellow) is buried in pocket B. (b) FabB$_{G107M}$=ACP$_{C8}$, FabF$_{wt}$=ACP$_{C8}$ in light blue (RMSD= 1.198), FabF$_{wt}$=ACP$_{C16}$ in pink (RMSD= 1.203), FabF$_{wt}$=ACP$_{C16}$ in teal (RMSD= 1.203), OXSM$_{wt}$=mACP$_{C8}$ in orange (RMSD= 1.651) are overlayed with FabB$_{G107M}$=ACP$_{C12}$. The overlay demonstrates that substrates have access to pocket A in FabB$_{wt}$, FabB$_{G107M}$, and FabF$_{wt}$. In FabF$_{wt}$, the opening of the gating loop (not shown) gives substrates access to both pocket A and the pocket behind the gate. The C16 substrate (teal) occupies pockets A and C. (c) A pocket surface from FabB$_{G107M}$ =ACP$_{C12}$. Truncation of pocket A by M107 redirects longer chains (i.e., C12) to pocket B. (d) A surface from FabB$_{wt}$ shows the C12 substrate in pocket A. (e) In FabF$_{wt}$=ACP$_{C8}$, the active site is more spacious than in FabB$_{wt}$, even with gate closed (Phe400 underneath the acyl chain) as a result of an A205 in place of M204 in FabB$_{wt}$. (f) In FabF$_{wt}$=ACP$_{C16}$, opening the Phe400 gate significantly increases the size of pocket C. When this gate opens, another pocket becomes accessible. The C16 substrates sample these two large pockets.

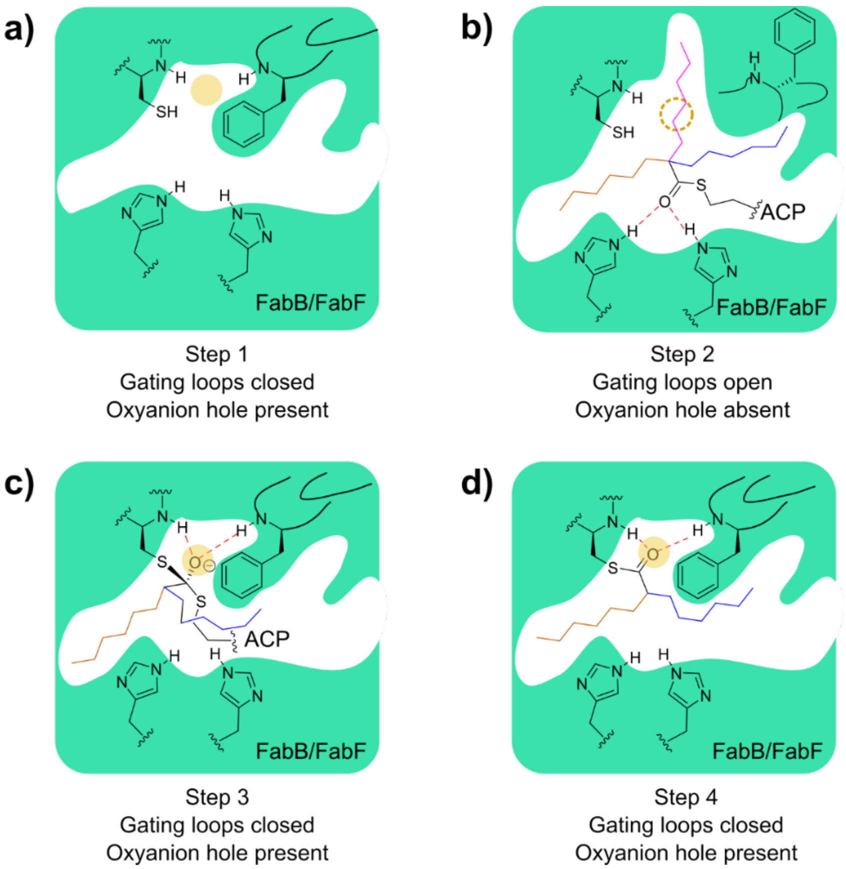


**Figure S19.** Transacylation steps in FabB or FabF. (a)Step 1 represents FabB or FabF in the *apo* form without substrate loaded. The oxyanion hole (yellow dot) is present while the gating loops are closed. Gating loop 1 contains a gating phenylalanine, while gating loop 2 does not. (b)In step 2, ACP delivers an acyl substrate into the binding pocket upon the opening of the gating loops. While the oxyanion hole is absent (dashed yellow circle), the substrate interacts with a pair of highly conserved His residues (His303 and His340 in FabF and His298 and His333 in FabB). During this state, the acyl chain has access to pocket A (brown), B (blue), and C (pink). (c)In Step 3, the gating loop closes to restore the oxyanion hole, which stabilizes the tetrahedral intermediate. In this state, which is mimicked by the crystal structures FabB$_{G107M}$=ACP$_{C8}$ and FabB$_{G107M}$=ACP$_{C12}$, the acyl chain samples pockets A and B. (d)Step 4 represents the end of the transacylation step, where the substrate is completely transferred onto the catalytic residue Cys. In this state, the acyl chain samples pockets A and B.

**SI TABLES**

**Table S1. Data collection and refinement statistics for X-ray crystallography**

| | $FabB_{G107M}$=$ACP_{C8}$ (PDB: 9MLW) | $FabB_{G107M}$=$ACP_{C12}$ (PDB: 9MLX) |
|---|---|---|
| **Data Collections** | | |
| Space group | P 1 21 1 | P 1 21 1 |
| Cell dimensions | | |
| *a*, *b*, *c* (Å) | 58.41, 98.18, 78.00 | 58.72, 98.25, 78.55 |
| α, β, γ (°) | 90.00, 109.25, 90.00 | 90.00, 109.32, 90.00 |
| Resolution (Å)[a] | 73.72 – 2.14 (9.07 – 2.14) | 59.17 – 2.25 (2.33 – 2.25) |
| $R_{merge}$[a] | 0.117 (0.617) | 0.076 (0.472) |
| $R_{pim}$[a] | 0.101 (0.535) | 0.176 (1.094) |
| $\langle I/\sigma(I) \rangle$[a] | 5.9 (1.9) | 9.2 (1.7) |
| CC1/2[a] | 0.99 (0.77) | 0.996 (0.757) |
| Completeness (%)[a] | 99.3 (94.4) | 99.9 (99.8) |
| Multiplicity[a] | 3.4 (3.1) | 3.3 (3.3) |
| **Refinement** | | |
| Resolution (Å) | 2.14 | 2.25 |
| No. reflections | 46000 | 56794 |
| $R_{work}$ / $R_{free}$ | 0.191/0.243 | 0.187/0.238 |
| No. atoms | | |
| Protein A, B, C, D | 3023, 3044, 637, 623 | 3022, 3050, 607, 593 |
| Water | 48 | 176 |
| *B*-factors | | |
| Protein A, B, C, D | 31, 32, 60, 52 | 29, 27, 61, 50 |
| R.m.s. deviations | | |
| Bond lengths (Å) | 0.007 | 0.62 |
| Bond angles (°) | 1.857 | 1.05 |

[a]Values in parentheses are for highest-resolution shell

**Table S2. Plasmids**

| Name | Plasmid Base | AntR * | ORI | Prom. * | Gene | Ind. * | Source |
|---|---|---|---|---|---|---|---|
| pET16b FabD | pET16b | Cb | pBR322 | $P_{T7}$ | *E. coli fabD* | IPTG | Ruppe et al.[2] |
| pET28a FabA | pET28A | Kan | pBR322 | $P_{T7}$ | *E. coli fabA* | IPTG | Ruppe et al.[2] |
| pET16b FabH | pET16b | Cb | pBR322 | $P_{T7}$ | *E. coli fabH* | IPTG | Ruppe et al.[2] |
| pET15b FabG | pET15b | Cb | pBR322 | $P_{T7}$ | *E. coli fabG* | IPTG | Ruppe et al.[2] |
| pET28a FabZ dimer | pET28A | Kan | pBR322 | $P_{T7}$ | *E. coli fabZ* | IPTG | Ruppe et al.[2] |
| pET16b FabI | pET16b | Cb | pBR322 | $P_{T7}$ | *E. coli fabI* | IPTG | Ruppe et al.[2] |
| pET28a FabB | pET28A | Kan | pBR322 | $P_{T7}$ | *E. coli fabB* | IPTG | Ruppe et al.[2] |
| pET16b FabF | pET16b | Cb | pBR322 | $P_{T7}$ | *E. coli fabF* | IPTG | Ruppe et al.[2] |
| pET28a TesA | pET28A | Kan | pBR322 | $P_{T7}$ | *E. coli ‘tesA* | IPTG | Andrzejewski et. al.[12] |
| pET22b ACP | pET22b | Cb | pBR322 | $P_{T7}$ | *E. coli ACP* | IPTG | Gift from Michael D. Burkart |
| pET28a FabB.G107M | pET28A | Kan | pBR322 | $P_{T7}$ | *E. coli fabB* | IPTG | Mains et. al.[13] |
